# The AAA+ ATPase RavA and its binding partner ViaA modulate *E. coli* aminoglycoside sensitivity through interaction with the inner membrane

**DOI:** 10.1101/2022.02.06.479298

**Authors:** Jan Felix, Ladislav Bumba, Clarissa Liesche, Angelique Fraudeau, Fabrice Rébeillé, Jessica Y. El Khoury, Karine Huard, Benoit Gallet, Christine Moriscot, Jean-Philippe Kleman, Yoan Duhoo, Matthew Jessop, Eaazhisai Kandiah, Frédéric Barras, Juliette Jouhet, Irina Gutsche

## Abstract

Enteric bacteria have to adapt to environmental stresses in the human gastrointestinal tract such as acid and nutrient stress, oxygen limitation and exposure to antibiotics. Membrane lipid composition has recently emerged as a key factor for stress adaptation. The *E. coli ravA-viaA* operon is essential for aminoglycoside bactericidal activity under anaerobiosis but its mechanism of action is unclear. Here we characterise the VWA domain-protein ViaA and its interaction with the AAA+ ATPase RavA, and find that both proteins localise at the inner cell membrane. We demonstrate that RavA and ViaA target specific phospholipids and subsequently identify their lipid-binding sites. We further show that mutations abolishing interaction with lipids restore induced changes in cell membrane morphology and lipid composition. Finally we reveal that these mutations render *E. coli* gentamicin-resistant under fumarate respiration conditions. Our work thus uncovers a *ravA-viaA*-based pathway which is mobilised in response to antibiotics under anaerobiosis and has a major impact on cell membrane regulation.

## Introduction

Although antibiotics are powerful antibacterial weapons, their widespread and frequently uncontrolled use has led to the emergence of numerous multi-resistant bacteria. Aminoglycosides (AGs) are highly efficient antibiotics against Gram-negative pathogens, yet they cause toxic side effects ^1^. Understanding what makes bacteria sensitive to AGs and why AG efficiency is notoriously reduced under anaerobic conditions, often encountered by intestinal pathogens, is essential since administration of smaller AG doses reduces their toxicity towards the host. In two paradigmatic enterobacteria, *Escherichia coli* and *Vibrio cholerae*, the expression of the *ravA-viaA* operon is enhanced by anaerobiosis and sensitises the bacteria to AGs, whereas a deletion of *ravA-viaA* enhances their AG resistance and reduces the AG-mediated toxic stress ^2–5^. The extensively studied ATPase RavA ^6–8^ belongs to the versatile AAA+ superfamily ^9,10^, whereas the largely uncharacterised ViaA carries a von Willebrand factor A (VWA) domain ^11^ and stimulates RavA ATPase activity ^6,12^. *E. coli* RavA-ViaA is suggested to act as a chaperone for the maturation of two respiratory complexes - Complex I (Nuo) ^3^ and fumarate reductase (Frd) ^12^. Increased AG uptake may therefore result from facilitated Nuo assembly that contributes to proton motive force generation. However, *E. coli* and *V. cholerae* respiratory pathways are very different, and *V. cholerae* lacks the Nuo complex ^13^. In addition, the documented RavA-ViaA-dependent inhibition of Frd activity ^12^ contradicts the hypothesis that *ravA-viaA* enhances respiration. Furthermore, *in vitro* RavA is sequestered by the acid stress-inducible lysine decarboxylase LdcI in the form of a huge LdcI-RavA cage proposed to function in the acid stress response upon oxygen and nutrient limitation ^6,8,14^. Thus, the molecular mechanism of RavA-ViaA action in bacterial physiology in general, and in AGs sensitisation in particular, appears enigmatic. Our recent study has unexpectedly revealed that LdcI forms supramolecular assemblies under the *E. coli* membrane, seemingly at lipid microdomains ^15^. Here we biochemically and biophysically characterise the ViaA protein and its interaction with RavA, and show that both proteins localise to the *E. coli* inner membrane, affect membrane morphology and modify cellular lipid homeostasis. We further reveal that both RavA and ViaA interact with specific lipids, identify the lipid-binding sites and demonstrate that their mutations abolish lipid binding *in vitro* and strongly attenuates the effect on lipid homeostasis *in vivo*. Moreover, we introduce these mutations into the *E. coli* chromosome and show that they abrogate the effect of the *ravA-viaA* operon on the AG sensitisation. Altogether, this work reveals a *ravA-viaA*-based pathway that is mobilised in the response of *E. coli* to AGs under anaerobic conditions and has a major impact on the regulation of the bacterial cell membrane. Building upon these results, we propose a possible scenario for the *ravA-viaA* function and open up further research perspectives.

## Results

### ViaA is a dimeric, soluble two-domain protein

The protein sequence of ViaA reveals that it has an N-terminal part (N-Terminal domain of ViaA, NTV) with a predominantly α-helical character that does not show sequence similarity to any other known protein (Figure 1A, residues 1 - 319). The C-terminal part (C-Terminal domain of ViaA, CTV), starting from residue 320, is predicted to be a VWA domain and contains the characteristic MIDAS motif (Figure 1A). While studies of the potential role of *E. coli* ViaA in the maturation of Nuo and Frd have been initiated ^3,12^, no protocols for the preparative purification of ViaA are available, its biochemical/biophysical characterization remains scarce, and binding to its potential interaction partners has only been shown indirectly ^3,6,12^. Addition of an N-terminal AviTag to the C-terminally His-tagged ViaA (hereafter named AviTag-ViaA-His) stabilised the protein and enabled its high yield purification for characterisation by multi-angle laser light scattering (MALLS) and small-angle X-ray scattering (SAXS) (Figure 1B-D, Materials and Methods). MALLS experiments with purified AviTag-ViaA-His sample showed a monodisperse peak and yielded a molecular weight (MW) of 123 kDa, corresponding to roughly two times the theoretical MW of the AviTag-ViaA-His construct (60.4 kDa). SAXS studies confirmed the dimeric state of ViaA in solution and resulted in predicted MWs of 130 and 117 kDa using SAXSMoW ^16,17^ and ScÅtter ^18^ respectively. Moreover, the distance distribution function calculated from the SAXS data suggests an elongated particle, while the Kratky plot reveals a degree of flexibility, and corresponds to a Kratky plot expected for a multidomain protein with flexible linker (Figure 1C, 1D). We note that the elongated character and apparent flexibility of ViaA might be accentuated by the presence of non-cleaved tags.

**Figure 1:**
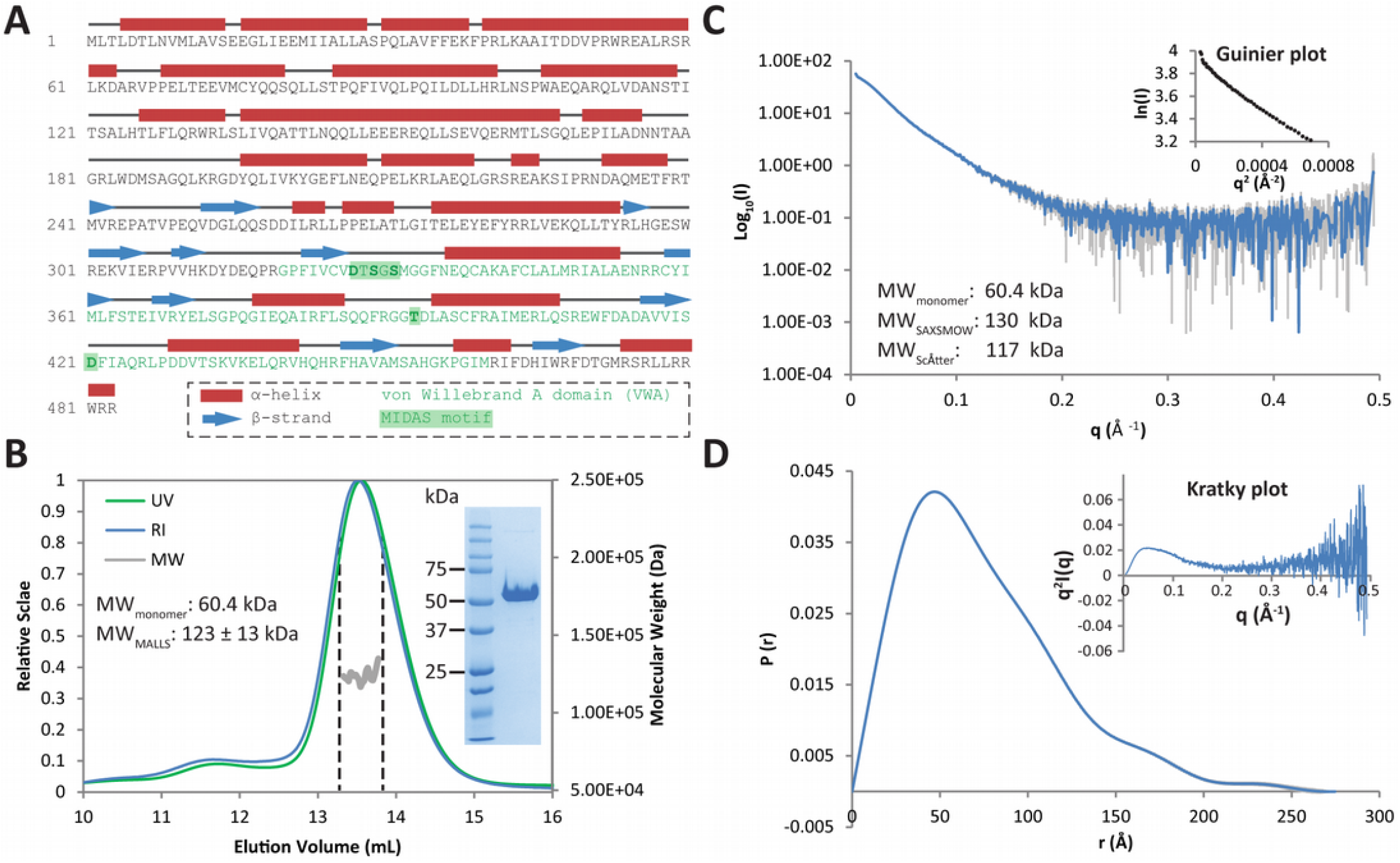
Biochemical and biophysical characterization of ViaA: A) Annotated sequence of *E. coli* ViaA, with predicted α-helices shown as red tubes and predicted β-strands shown as blue arrows. The von Willebrand A (VWA) domain of ViaA is shown in green, with the MIDAS motif highlighted in dark green. B) Molecular mass determination of AviTag-ViaA-His by SEC-MALLS. The differential refractive index (RI) signal is plotted (left axis, blue curve) along with the UV signal (UV, left axis, green curve) and the determined molecular weight (MW, right axis, grey curve). An SDS-PAGE gel of purified AviTag-ViaA-His is shown as an inset on the right-hand side of the plot. The theoretical monomer MW and the MW as determined by MALLS are annotated on the left-hand side of the plot. C) SAXS curve of purified AviTag-ViaA-His. An inset shows a Guinier Plot zooming in on the low-q region of the scattering curve. The theoretical monomer MW as well as the MW determined by SAXSMOW and ScÅtter are shown on the left. D) Pair-wise distance distribution function *P(r)* and Kratky plot (Top right inset) of the AviTag-ViaA-His SAXS data.

### The strength and kinetics of the RavA-ViaA interaction are nucleotide-dependent

While a previous observation that ViaA or NTV slightly enhance the ATPase activity of RavA suggested a physical interaction between the two proteins via the N-terminal part of ViaA ^6,12^, direct measurements of the RavA-ViaA interaction are unavailable. Pull-down assays using SPA-tagged RavA did not bring down ViaA, and vice versa, pointing to a rather weak or transient interaction ^12^. The advantage of AviTag-containing constructs is that they can be biotinylated *in vitro* and coupled to Streptavidin (SA)-biosensors for binding studies using Bio-Layer Interferometry (BLI). To address potential effects of the protein orientation on the biosensor, either N- or C-terminally tagged constructs (Materials and Methods) were immobilised and transferred to wells containing a concentration series of purified RavA in the absence or presence of ADP or ATP (Figure 2A, 2B). The affinity of the apo-RavA to immobilised His-ViaA-AviTag and AviTag-ViaA-His could be described by dissociation constants (K_D_) of 11.0 µM and 18.9 µM respectively, showing that the location of the AviTag does not have a major influence and validating the weak nature of the RavA-ViaA interaction. ADP binding to RavA increased the strength of the RavA-ViaA interaction to ∼2.2 µM, whereas the presence of ATP altered the shape of the BLI curves that could only be reliably fitted using a 2:1 heterogeneous ligand binding model, resulting in K_D1_/K_D2_ values of 30.3/1.0 µM and 37.5/0.5 µM for His-ViaA-AviTag and AviTag-ViaA-His respectively. Taken together, these BLI binding studies revealed that ViaA directly interacts with RavA, and that the strength of this interaction is dependent on the nucleotide-bound state of RavA. Complementary BLI studies using NTV-AviTag demonstrated similar RavA binding kinetics as the full-length ViaA despite the monomericity of the NTV construct (Supplementary Figure 1), confirming that NTV is sufficient for RavA binding ^12^. In addition, the absence of the LARA domain of RavA responsible for the LdcI-RavA cage formation ^7,14,19^ did not affect the binding of RavA to ViaA (Supplementary Figure 1), indicating that the LARA domain is not involved in the RavA-ViaA interaction.

**Figure 2:**
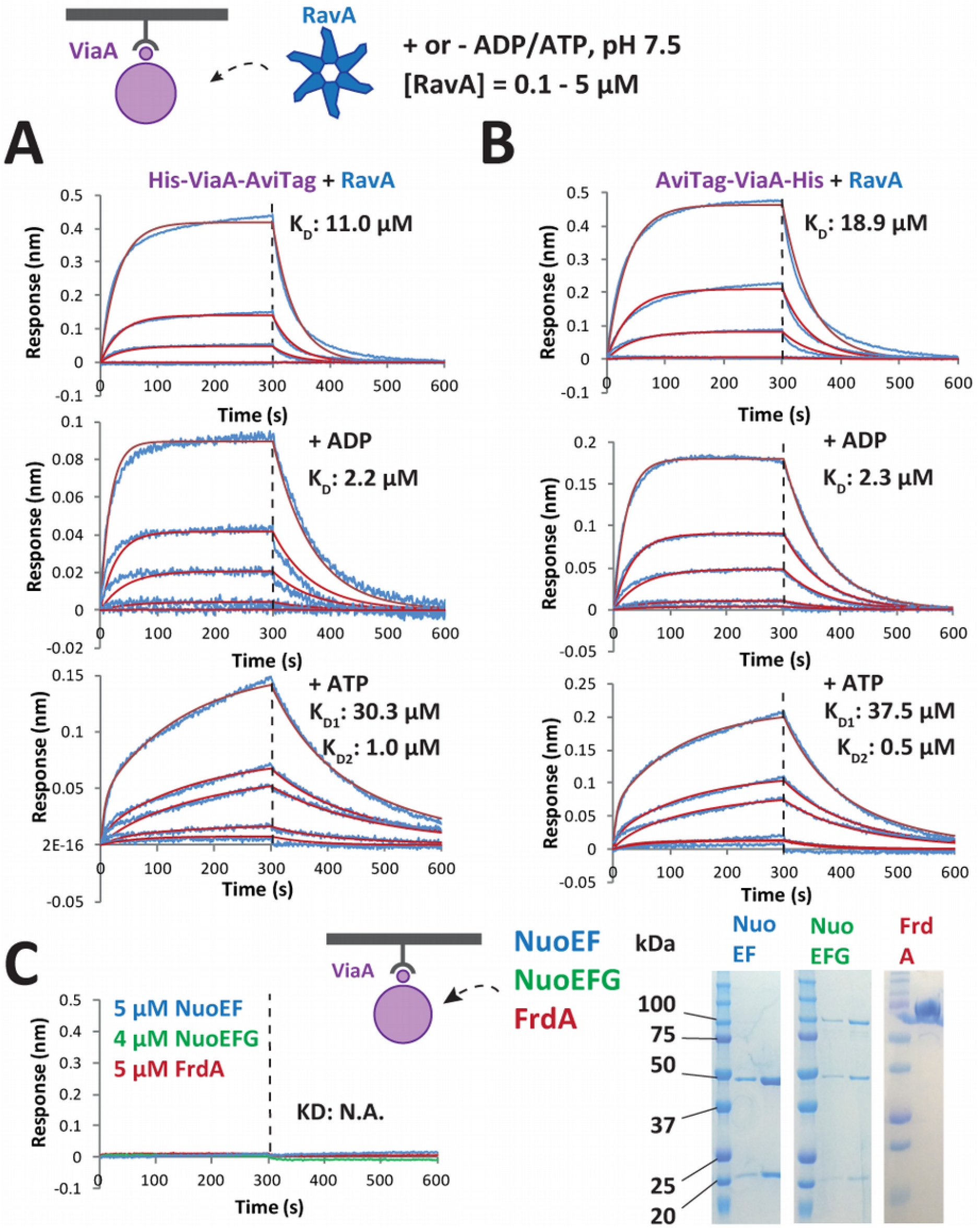
Characterization of the ViaA-RavA interaction by Bio-Layer Interferometry (BLI) binding studies. A & B) BLI measurements of His-ViaA-AviTag (A) or AviTag-ViaA-His (B) coupled on BLI biosensors and RavA, with or without added ADP/ATP. The blue curves correspond to the measured signal while the red curves correspond to the calculated fit using a 1:1 (no nucleotide, ADP) or 2:1 heterogeneous ligand binding (ATP) interaction model. C) BLI measurements of AviTAg-ViaA-His coupled on BLI biosensors and either NuoEF (blue curve), NuoEFG (green curve) or FrdA (red curve). SDS-PAGE gels of purified NuoEF, NuoEFG and FrdA are shown on the right.

Previous observation that upon cell fractionation RavA was mostly present in the cytoplasmic fraction whereas the majority of ViaA partitioned into the inner membrane fraction ^3^, led the authors to the detection of ViaA interaction with specific subunits of Nuo ^3^ and Frd ^12^ by pull down assays with SPA-tagged baits. Thus, using BLI, we interrogated the interaction between ViaA and its proposed respiratory complexes’ partners: the soluble catalytic NADH dehydrogenase domain of *E. coli* NADH:Ubiquinone Oxidoreductase I (NuoEFG) and the flavin-containing subunit of the *E. coli* Fumarate Reductase (FrdA) ^3,12^. Expression and purification of NuoEFG and FrdA were performed following previously published protocols ^20–22^ (Materials and Methods), and resulted in purified fractions with a red/brown and a yellow color for NuoEFG and FrdA respectively, corresponding to the presence of Riboflavin/Iron-Sulfur clusters for NuoEFG and FAD for FrdA. However, using concentrations as high as 5 µM of NuoEFG and FrdA did not result in any discernible interaction with either His-ViaA-AviTag or AviTag-ViaA-His (Figure 2C).

### Both RavA and ViaA show membrane localisation in *E. coli* cells

Surprised by the absence of *in vitro* interactions between ViaA and its purified proposed binding partners, we wondered what might then be the reason for its described partitioning into the inner membrane fraction ^3^, and therefore sought to localise ViaA and RavA *in cellulo* by single molecule localisation microscopy imaging. Considering the low amount of natively expressed RavA and ViaA ^3^, we opted for overexpression (OE) of these proteins with particular tags (collectively called RavA-OE and ViaA-OE for overexpressed RavA and ViaA constructs respectively). The N- or the C-terminus of RavA was tagged with a 12 amino acid BC2 peptide and the resulting RavA-OE was immunolabelled with an anti-BC2 nanobody ^23^ coupled to the Alexa Fluor 647 fluorescent dye for stochastic optical reconstruction microscopy (STORM) imaging. ViaA-OE constructs were produced by fusion of the PAmCherry fluorescent protein ^24^ to either the N- or the C-terminus of ViaA, and imaged by photo-activated localisation microscopy (PALM). In addition, PAmCherry was fused to the C-terminus of the NTV and to the N-terminus of the CTV to make NTV-OE and CTV-OE respectively.

Unexpectedly, although RavA is presumed to be cytoplasmic, RavA-OE showed a propensity to localise at the cell periphery rather than being distributed homogeneously through the volume (Figure3A). Furthermore, both ViaA-OE constructs, as well as CTV-OE, also exhibited a distinct peripheral localisation pattern around the entire circumference of the cell, with a tendency to accumulate at the polar regions (Figure 3A). The NTV-OE, in contrast, formed cytosolic bodies (Supplementary Figure 2). Wide-field images of His-RavA and His-ViaA overexpressing *E. coli* cells labelled with specific anti-RavA and anti-ViaA nanobodies coupled to Alexa Fluor dyes (Materials and Methods) further confirmed the inner membrane distribution of RavA and ViaA (Supplementary Figure 2). Altogether, these experiments indicate that *in cellulo* ViaA is targeted to the bacterial inner membrane despite the absence of ViaA binding to purified NuoEFG and FrdA *in vitro*, with the CTV being required for this targeting.

**Figure 3:**
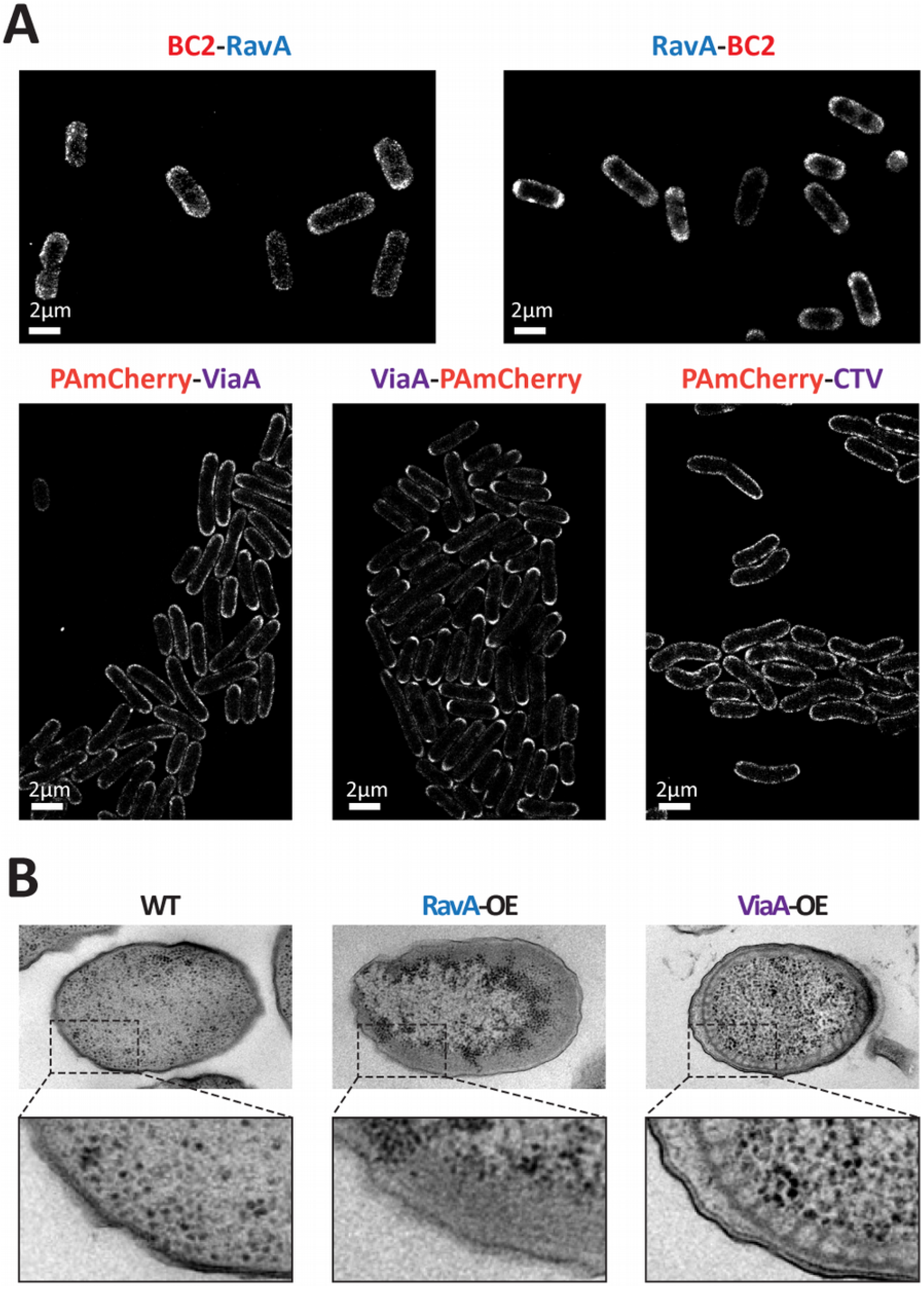
Imaging of overexpressed RavA and ViaA by super-resolution microscopy and cellular TEM. A) Single molecule localisation microscopy imaging of *E. coli* cells overexpressing fluorescently labeled RavA and ViaA by STORM (BC2-RAvA, RavA-BC2) and PALM (PAmCherry-ViaA, ViaA-PAmCherry and PAmCherry-CTV). B) Imaging of high pressure frozen, freeze-substituted and sectioned wild-type (WT) *E. coli* cells and *E. coli* cells overexpressing RavA (RavA-OE) or ViaA (ViaA-OE) by TEM.

### RavA accumulates under the inner membrane whereas ViaA modifies membrane morphology of *E. coli* cells

In parallel to imaging the RavA-OE and ViaA-OE bacteria by epifluorescence and super-resolution microscopy, these bacteria were subjected to high pressure freezing and freeze substitution followed by ultrathin cell sectioning, and their ultrastructure examined by transmission electron microscopy (EM). A dark layer of matter, supposedly corresponding to the RavA protein, accumulated under the bacterial inner membrane, in line with the fluorescence microscopy observations (Figure 3B). Most strikingly, arrays of ectopic intracellular membrane tubes were discovered to underlie the inner membrane of the ViaA-OE bacteria (Figure 3B). Interestingly, overexpression of few membrane proteins with particular topologies has previously been noticed to induce formation of tubular membrane networks of reminiscent hexagonal phase morphologies ^25,26^. Such cardiolipin (CL)-enriched neo-membranes are proposed to form in order to accommodate the highly overproduced membrane protein, thereby relieving the inner cellular membrane from the associated stress. Thus, on the one side we have biochemically and biophysically characterised a soluble form of ViaA *in vitro*, and on the other side our morphological cellular EM observations agree with and corroborate our optical imaging data that show localisation of ViaA and RavA at the inner membrane of the bacterial cell. This raises two immediate questions: (i) are ViaA and RavA capable of directly binding to lipids, and (ii) do they modulate cellular lipid homeostasis, and in particular, does ViaA overexpression increase the cellular amounts of CL? In this respect, it is interesting to note that the conical shape of CL is known to favor its clustering in negative curvature regions such as cell poles and the division septum ^27,28^, which would agree with the localization of the overexpressed ViaA that we observed by PALM.

### Both RavA and ViaA bind specific anionic phospholipids *in vitro*

The inner membranes of *E. coli* consist of three major anionic phospholipids: 70-80% phosphatidylethanolamine (PE), 20-25% phosphatidylglycerol (PG) and ∼5% cardiolipin (CL), the exact content of the latter being culture condition and growth phase-dependent ^29–31^. The universal precursor of these phospholipids is phosphatidic acid (PA). Our dot-blot assays (Figure 4A, Supplementary Figure 3) revealed that purified RavA specifically binds PG, whereas ViaA specifically binds PA and to a lesser extent CL. RavA interactions with ViaA and PG are mutually exclusive; likewise, inside the LdcI-RavA complex, RavA is no longer able to interact with PG. In contrast, lipid-bound ViaA is still capable of interacting with RavA (Figure 4A, Supplementary Figure 3). Surprisingly, RavA_ΔLARA_ loses the PG binding capacity (Supplementary Figure 3), which shows that the LARA domain is implicated not only in the interaction with LdcI but also in the interaction with lipids. In this light, and since the LARA domain is involved in lipid binding but not in ViaA binding (Supplementary Figure 1), the observation that RavA cannot simultaneously interact with ViaA and PG suggests that binding to PG may induce a conformational change in the RavA molecule, making it inapt for ViaA binding.

**Figure 4:**
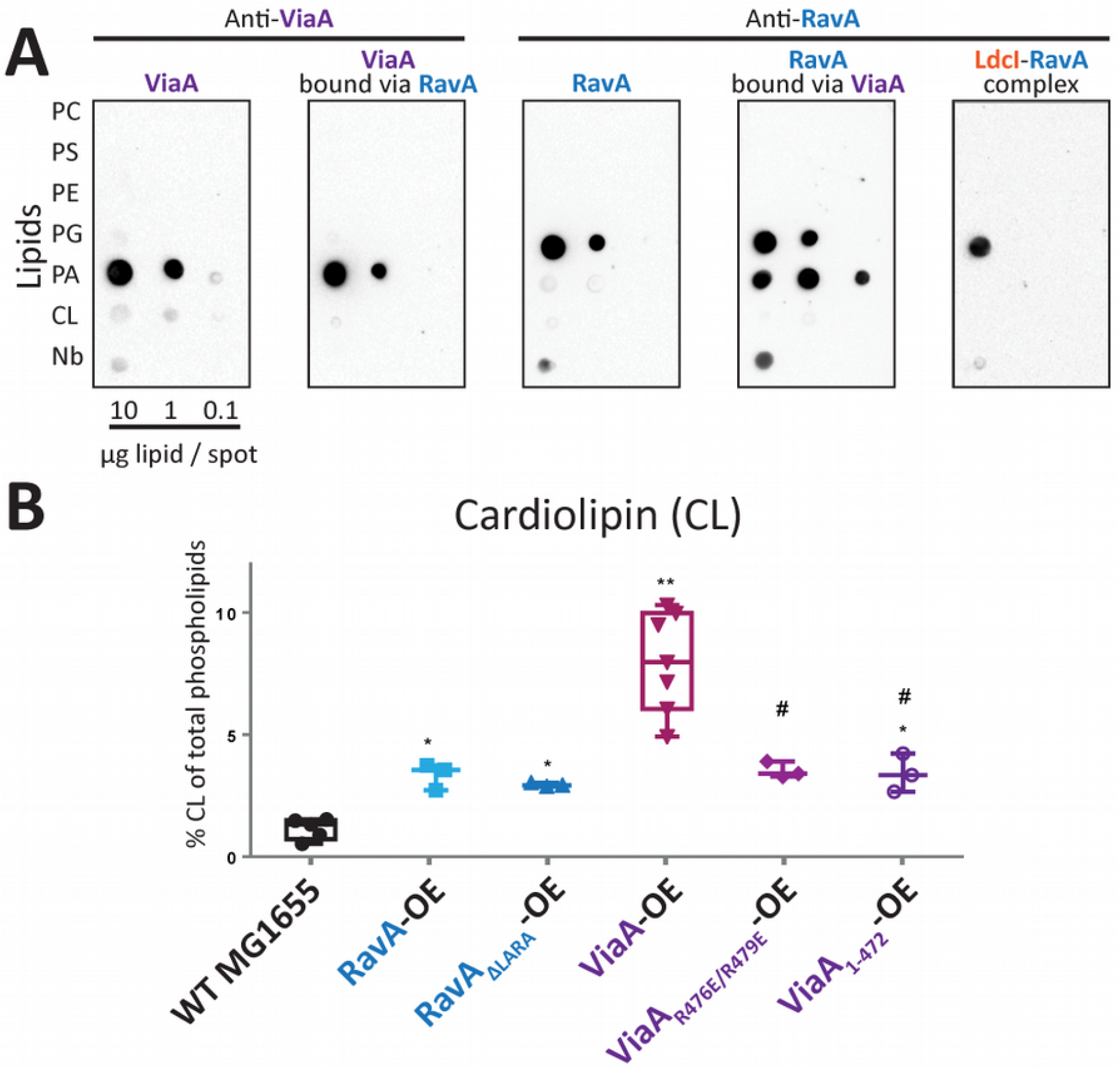
RavA and ViaA binding to specific lipids, and quantification of CL in different RavA and ViaA overexpressing strains. A) Dot-blot assays using purified ViaA and RavA, visualized using anti-ViaA and anti-RavA antibodies and a secondary HRP-antibody conjugate (PC: phosphatidylcholine, PS: phosphatidylserine, PE: phosphatidylethanolamine, PG: phosphatidylglycerol, PA: phosphatidic acid, CL: cardiolipin). For ‘ViaA bound via RavA’ or ‘RavA bound via ViaA’: Lipid binding was first allowed for RavA and subsequently ViaA was added, or vice versa. For ‘LdcI-RavA-complex’, proteins were mixed before incubation on the membrane (see also Materials and Methods). B) Quantification of cardiolipin (CL) levels by TLC and GC-FID/MS in wild-type (WT) MG1655 *E. coli* cells and MG1655 *E. coli* cells overexpressing RavA, RavA_ΔLARA_, ViaA, ViaA_R476E/R479E_ and ViaA_1-472_, visualized by a Tukey representation (showing all points from min. to max.). Each point represents a biological repeat. The overexpressing (OE) series were compared with the control WT MG1655 series using an unpaired nonparametric Mann-Whitney test. A significant difference with the control is shown by * (p value < 0.05) or ** (p value < 0.01). The # indicates a significant difference (p value < 0.05) with ViaA OE.

The observation that ViaA can simultaneously interact with RavA and with lipids is consistent with ViaA being a two-domain protein, with the NTV binding to RavA and the CTV binding to lipids. Since the 3D structure of ViaA is unknown, we examined its primary sequence and the recent 3D structure prediction of the CTV provided by Alphafold ^32,33^ and RoseTTAFold ^34^ (Figure 5) for hints to possible positions of a lipid binding site. The C-terminal end of ViaA is highly positively charged, with eight arginines, six of which are situated in the predicted α-helix at the C-terminal extremity (Figure 5A). Compelled by the amphipatic nature of this helix (Figure 5B) and its high conservation among enterobacterial ViaA proteins (Figure 5C), we reasoned that the C-terminus of ViaA might be involved in electrostatic interactions with the negatively charged head groups of PA and CL. In particular, R476 and R479 are predicted to be situated at the same side of the α-helix and seem to form a positive patch (Figure 5B). To evaluate this hypothesis, we designed the three following mutants: ViaA_1-472_, devoid of the entire C-terminus, ViaA_R476A/R479A_ and ViaA_R476E/R479E_. As shown in Supplementary Figure 3, mutations of R476 and R479 into alanines notably attenuate interaction with lipids, whereas the removal of the C-terminus and the R-to-E double mutation result in the abrogation of the lipid-binding propensity. Altogether, combined with the membrane distribution and effects on membrane morphology, ViaA unexpectedly joins the list of peripheral membrane proteins tethered to the inner *E. coli* membrane through an amphipathic helix displaying affinity for anionic lipids and in particular CL, which also includes proteins such as the cell division site selection ATPase MinD ^35,36^ and the cell shape-determining actin homolog MreB ^37^. Strikingly, although ViaA is a peripheral membrane protein, its membrane-targeting C-terminal sequence contains all the key features of a high-affinity CL-binding site on bacterial integral membrane proteins, recently identified by molecular dynamics simulations ^38^: two or three adjacent basic residues, at least one polar, a neighbouring aromatics and a glycine proposed to confer a higher affinity for the CL head group.

**Figure 5:**
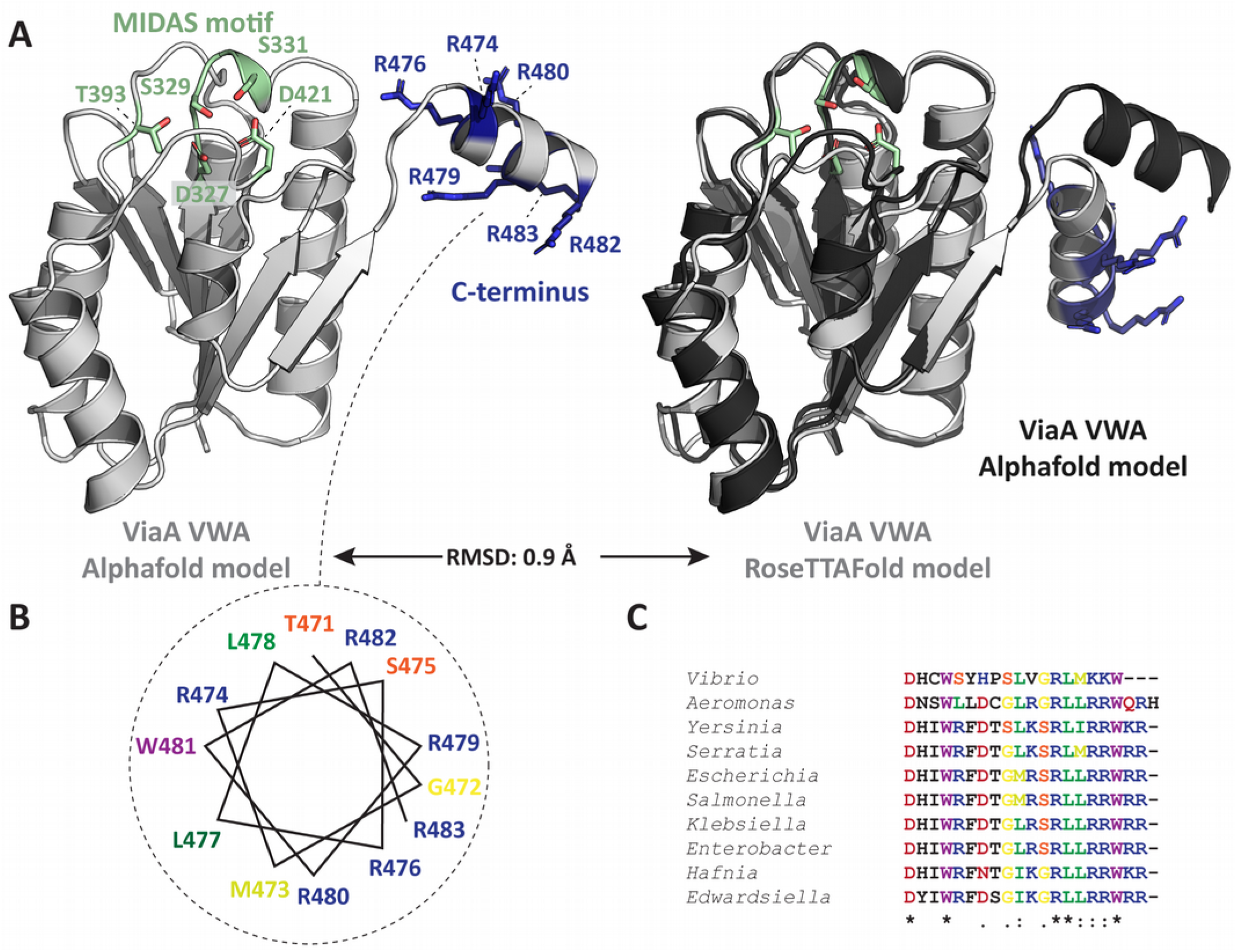
A) Cartoon presentation of Alphafold (left, light grey) and RoseTTAFold (right, light grey) structural predictions for the ViaA VWA domain. The MIDAS motif (green) as well as Arginine residues in the C-terminal helix (dark blue) are annotated and shown as sticks. The Alphafold and RoseTTAFold models are shown superimposed on the right in dark and light grey respectively. B) Helical wheel diagram of the C-terminal helix of ViaA (residues 471 – 483), with annotated and color-coded residues. C) Multiple sequence alignment of the C-terminal helix of different enterobacterial ViaA orthologues. Residues are color-coded and level of conservation is annotated with ‘*’ (perfect alignment), ‘:’ (strong similarity) and ‘.’ (weak similarity).

Finally, to further validate the proposed lipid binding sites of RavA and ViaA, we examined the morphology of RavA_ΔLARA_-OE, ViaA_R476E/R479E_-OE and ViaA_1-472_ -OE cells by EM and compared it to the observations presented in Figure 3B. Remarkably, these mutants formed soluble cytosolic bodies inside the bacterial cells (Supplementary Figure 4), although the overexpression levels remained very similar to those of the lipid binding-competent counterparts (Supplementary Figure 5). Thus, the inner membrane localisation and the morphological changes observed with RavA-OE and ViaA-OE are unequivocally related to their propensity to target specific phospholipids.

### ViaA modifies cellular lipid homeostasis *in vivo*

We then investigated the effect of RavA or ViaA overexpression on the membrane lipid composition (Figure 4B, Supplementary Figure 6). To this end, lipids were extracted from different cellular samples, and analysed by gas chromatography, thin layer chromatography and mass spectrometry (see Methods). As shown in Supplementary Figure 7, RavA-OE, RavA_ΔLARA_-OE, ViaA_R476E/R479E_-OE and ViaA_1-472_ -OE bacteria contained slightly but statistically significantly fewer lipids, quantified by their fatty acids, than the wild type (WT) MG1655 cells, whereas the total fatty acid content in ViaA-OE remained roughly similar to the WT. A similar behaviour was observed for the relative proportion of PG among the total lipids, whereas the relative proportion of PE remained unchanged. As expected from the electron microscopy observations, the most spectacular changes were observed with CL, the proportion of which was 6 to 10-fold increased in the ViaA-OE cells, whereas overexpression of the other constructs led to a much milder (but still statistically significant) CL increase in comparison to the WT bacteria (Figure 4B). The low native expression level of ViaA precludes conclusive assessments of the potential subtle difference in the lipid metabolism between the WT and the MG1655-ΔravAviaA strains under anaerobic conditions favoring expression of the *ravA-viaA* operon. One should however keep in mind that as observed for some rare membrane proteins, an inner membrane stress due to its saturation resulting from the ViaA overexpression may be relieved by formation of the CL-enriched neo-membranes that enclose the excess of ViaA.

**Figure 6:**
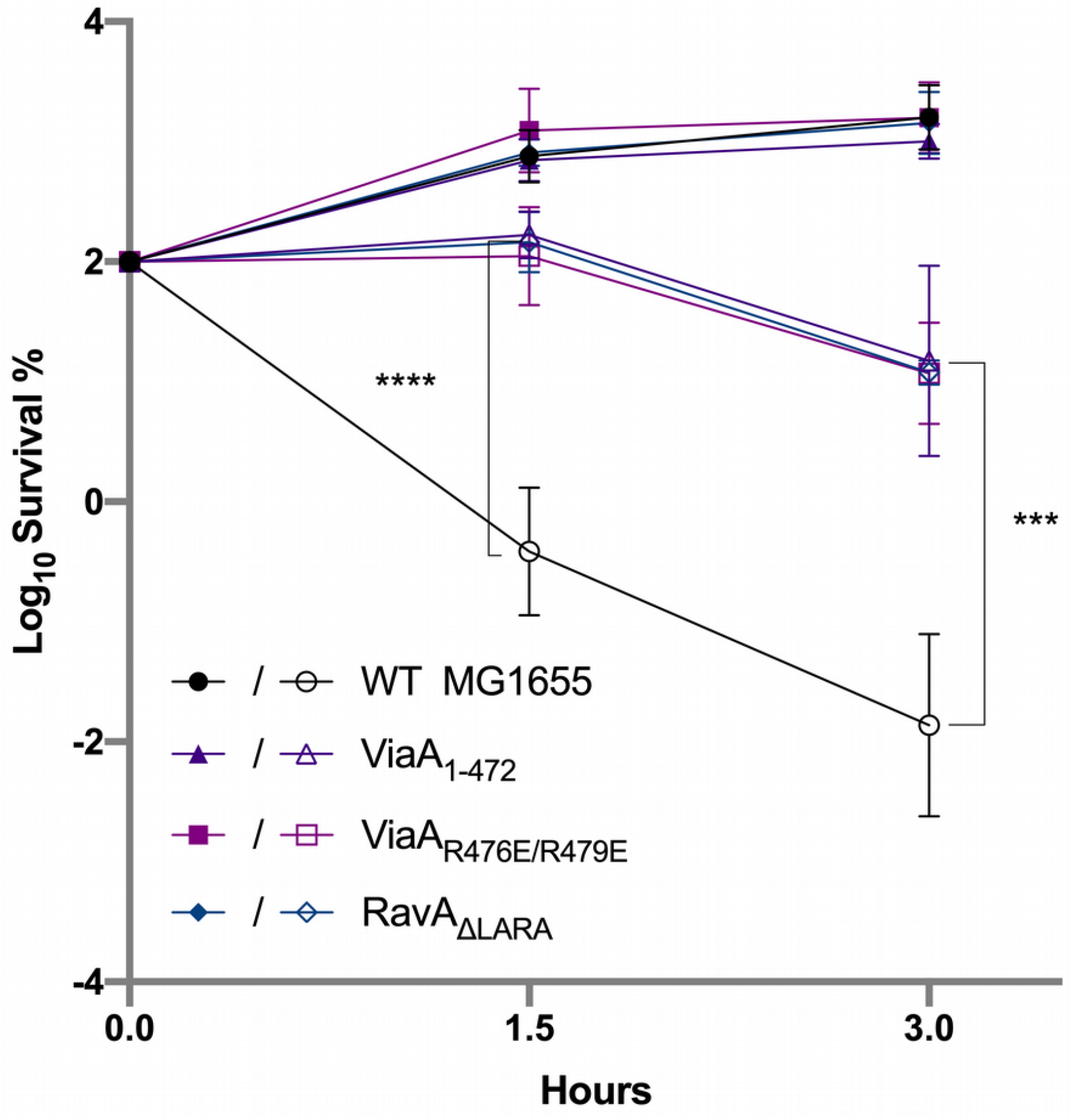
Lipid binding-deficient chromosomal mutants of RavA and ViaA impair RavA and ViaA function in gentamicin sensitisation. Survival of wild type MG1655, ViaA_1-472_, ViaA_R476E/R479E_ and RavA_*ΔLARA*_ *E. coli* strains after Gm treatment. Cells were grown to an OD_600_ of 0.2 in LB supplemented with 10 mM fumarate, after which 16 µg/mL Gm was added. The survival values after 1.5 and 3 hours of treatment are represented. Full and empty symbols are for untreated and Gm-treated bacteria, respectively. Survival, measured by colony-forming units (CFU) per mL, was normalized relative to time zero at which Gm was added (early log phase cells; ∼5×10^7^ CFU/mL) and plotted as Log_10_ of % survival. Values are expressed as a mean value of 3 biological replicates and error bars depict standard deviations. A one-way ANOVA test followed by a Dunnett’s multiple comparisons test was performed to compare at each time point (1.5 & 3 hours) the treated WT to each of the treated mutant strains (*** adjusted *p*Value = 0.0002 & **** adjusted *p*Value < 0.0001).

### Lipid binding-deficient chromosomal mutants of RavA and ViaA impair RavA and ViaA function in gentamicin sensitisation

Having recently shown that *ravA-viaA* sensitise *E. coli* to gentamicin (Gm) in fumarate respiratory condition ^5^, we decided to check if lipid binding-deficient chromosomal MG1655 mutants ViaA_1-472_, ViaA_R476E/R479E_ and RavA_ΔLARA_ are still capable of Gm sensitisation. To this end, we performed a time-dependent killing experiment, using a Gm concentration equivalent to twice the minimum inhibitory concentration. As shown in Figure 6, all mutants exhibited increased resistance to Gm compared to the WT strain. This result demonstrates that RavA and ViaA mutants, designed to be unable to bind lipids, prevent Gm toxicity under fumarate respiratory conditions.

An attractive hypothesis could be that ViaA, eventually with the help of RavA, would sensitise *E. coli* to Gm by directly binding to the membrane and modifying its permeability. To address this possibility, we performed planar lipid bilayer experiments (Supplementary Figure 7) that confirmed binding of single molecules of RavA and ViaA to membranes *in vitro* but did not detect formation of membrane pores that would facilitate AG transport. Therefore, we concluded that RavA and ViaA target phospholipids without perforating the lipid bilayer such as to directly enable Gm uptake into the bacterial cell.

## Discussion

Since AGs are notoriously inefficient under the anaerobic and acidic conditions of the human gastrointestinal tract, insights into the mechanism of the *ravA-viaA* function in *E. coli* sensitisation to AGs should improve our understanding and fight against the global threat of antibiotic resistance. The *E. coli* MoxR AAA+ ATPase RavA has long been known to tightly bind one of the main enterobacterial acid stress response proteins, the acid stress inducible lysine decarboxylase LdcI, thereby preventing its inhibition by the nutrient stress response alarmone ppGpp ^6,7^. However, a characteristic of the MoxR AAA+ ATPases is also the close co-occurence of their genes with genes encoding VWA domain-containing proteins, with which they are functionally linked ^6^. The *ravA-viaA* operon, shown to be controlled by the anaerobic transcriptional regulator Fnr ^12^, sensitises *E. coli* to AGs in respiratory conditions when fumarate is used as electron acceptor instead of O_2_ ^5,39^. Thus, we decided to focus this work on a thorough characterisation of ViaA and the RavA-ViaA interaction, and investigated their molecular functions beyond the LdcI-RavA interaction.

We now provide extensive *in vitro* and *in vivo* biochemical, biophysical, optical imaging and morphological evidence that RavA and ViaA do not only interact in a nucleotide-dependent fashion, but also localise at the bacterial inner membrane, bind specific phospholipids and modulate membrane lipid composition. In particular, we demonstrate specific RavA-PG and ViaA-PA/CL interaction, and reveal a prominent effect of ViaA on the CL-enrichment of the *E. coli* inner membrane. A possible explanation of the effect of ViaA on CL can be proposed based on our finding that ViaA interacts strongly with PA (Figure 4A). Indeed, PA, PG and CL are tightly interconnected: PA, only present in tiny amounts in *E. coli* cells, is a metabolic hub for PG synthesis, whereas two PG molecules are necessary to produce CL ^40^. The synthesis of CL in bacteria requires that a first PG receives a phosphatidyl group from a second PG by a transesterification reaction catalysed by cardiolipin synthase (Cls). *E. coli* contains three Cls isoforms ^41^, with ClsA being the major contributor to CL synthesis during exponential growth. Interestingly, PA was shown to strongly inhibit ClsA activity ^42^. Consequently, the association of ViaA with PA at the inner membrane level could trap PA, thereby preventing Cls inhibition and eventually favouring CL synthesis, leading to a modification in the lipid homeostasis. Independently of the concrete mechanism, one may anticipate that by binding to PA and CL, natively expressed ViaA would influence some local membrane properties, for example by acting on the local membrane curvature, affecting membrane permeability or coming into close proximity to proteins present in CL-enriched lipid microdomains. Excitingly, such microdomains form CL-based proton sinks ^43^ that compartmentalise oxidative phosphorylation complexes ^44,45^ such as Nuo and Frd and are proposed to also attract LdcI ^15^.

Importantly, we show that the RavA and ViaA lipid-binding propensity is directly linked to their effect on the AG bactericidal activity under anaerobiosis, because mutations abolishing interaction with lipids preclude Gm toxicity under conditions of fumarate respiration. Our finding that RavA and ViaA do not permeabilise the lipid bilayer, and the independent observations by us and by others that anaerobic RavA-ViaA-mediated AG sensitivity is dependent on the proton motive force (pmf) ^3,5,39^, suggest that they may positively act on Frd or other respiratory complexes, thereby enhancing the pmf required for efficient AG uptake. Considering the absence of a direct interaction with purified FrdA and NuoEFG in a BLI setup as described above, as well as the lipid binding and membrane remodelling activity of RavA and ViaA revealed in the present study, we propose that RavA and ViaA chaperone certain respiratory complexes by acting on lipid microdomains in which these complexes are inserted. This hypothesis aligns with our recent observation of a patchy peripheral distribution of LdcI, which we attributed to an attraction of this proton-consuming acid stress response enzyme to anionic phospholipids domains forming proton sinks that compartmentalise oxidative phosphorylation complexes ^15^. Thus, this work sets the stage for future investigations of an unprecedented molecular network that links the LdcI-RavA-ViaA triad with bacterial stress adaptation, membrane homeostasis, respiratory complexes and aminoglycoside bactericidal activity.

## Materials and Methods

### Bacterial strains and plasmids construction

All plasmids, primers, and cloning strategies are summarized in Supplementary Table 1. The *ravA* and *viaA* genes were PCR amplified from *Escherichia coli* K-12 MG1655 genomic DNA using Phusion polymerase (Biolabs). All PCR products were purified by DNA cleanup kit (Qiagen). Gibson assemblies were performed using 0.4 U T5 exonuclease, 2.5 U Phusion polymerase, and 400 U Taq ligase (New England Biolabs) in 1× ISO buffer consisting of 100 mM Tris·HCl pH 7.5, 10 mM MgCl_2_, 0.8 mM dNTP mix, 10 mM dithiothreitol (DTT), 50 mg polyethylene glycol (PEG)-8000, 1 mM nicotinamide adenine dinucleotide (NAD). A total of 7.5 µL of the Gibson Master Mix was mixed with 2.5 µL DNA, containing ∼100 ng of vector. The mix was incubated for 60 min at 50 °C. Transformations were performed in Top10 competent bacteria (One Shot TOP10 chemically competent *E. coli*, Invitrogen) and selected using 100 µg/mL ampicillin or/and 34 µg/mL chloramphenicol (Euromedex). Agarose gel purification and DNA plasmid extractions were performed using a QIAquick Gel extraction kit (QIAGEN). Restriction enzyme digests were performed according to manufacturer recommendations (Biolabs). Bacteria were made electrocompetent by growing cells to an OD_600_ of around 0.7. Cells were then placed on ice and washed three times with ice-cold, sterile water. Electroporation using the MicroPulser (BioRad) was followed by 1 h growth in fresh LB medium.

Mutant strains of K-12 substrain MG1655 (ATCC 47076) were generated by recombineering following the protocols of earlier publications ^46,47^. Briefly, the pKD46 plasmid was used to express the three proteins Gam, Exo, and Beta, that mediate homologous recombination (HR) and that are originally derived from the bacteriophage lambda. On pKD46, expression of those genes is controlled by the *araBAD* promoter. pKD46 has an ampicillin resistance gene and a temperature sensitive origin of replication (ORI), hence cells with pKD46 were grown at 30°C with ampicillin. Recombineering requires DNA substrates with regions homologous to the target genome. Those substrates were produced by PCR using specifically designed pKD3-plasmids serving as PCR-template. pKD3 encodes the gene *CAT* allowing selection of engineered bacteria using chloramphenicol. The *CAT* cassette is flanked by FRT sites and can be removed using FLP recombinase. FLP is expressed from the pCP20 plasmid. pCP20 has a temperature-sensitive ORI and encodes the ampicillin resistance gene *ampR*, hence cells were grown at 30°C with ampicillin to keep the plasmid. To induce expression of FLP and to lose the plasmid, cells were grown at 43°C.

To engineer the here described mutant strains MG1655 ΔravA/viaA(1-472)::cat, MG1655 ΔravA/viaA-R476E/R479E::cat and MG1655 ravAΔLARA/ΔviaA::cat by HR, we targeted the following genomic sequence flanking the *ravA/viaA* operon: (5’-GTATGGCCAGCTGCTGTTCGCGAGAGCGTCCCTTCTCTGCTGTAAGCCATGGTCCATATGAATATCC-3’) and (5’-CTCGCAATTTACGCAGAACTTTTGACGAAAGGACGCCACTTCATTATGGCTCACCCTCATTTATTAGC-3’). To obtain PCR products for HR that encode mutant *viaA* and lack *ravA*, the corresponding mutant *viaA* genes were cloned into pKD3 (pKD3_ViaA(1-472), pKD3_ViaA(R476E/R479E)). Likewise, to get PCR products for HR that encode mutant *ravA* and lack *via*A, the corresponding mutant *ravA* gene was cloned into pKD3 (pKD3_RavA-ΔLARA).

For recombineering, the pKD46 plasmid was electroporated into MG1655ΔravA/ΔviaA target cells and grown on ampicillin plates over night at 30°C. On the second day, single colonies of MG1655ΔravA/ΔviaA:pKD46 were re-streaked on ampicillin plates and grown overnight at 30ºC. In the afternoon of the third day, a 20 ml pre-culture of MG1655:pKD46 cells was prepared and grown over night at 30ºC. On the fourth day, in order to induce expression of *gam, exo* and *beta*, about 3 ml of pre-culture was diluted into 100 ml fresh medium containing ampicillin and 0.2% arabinose (final concentration) and grown to an optical density of about 0.7. Cells were immediately made electrocompetent and electroporated with the specific PCR product that served as template for homologous recombination. All cells from the electroporation cuvette were plated on chloramphenicol plates and grown overnight at 37°C. On the fifth day, (all) single clones were re-streaked on fresh chloramphenicol plates for overnight growth at 37°C. The next day, single colonies were tested by colony PCR and re-streaked on chloramphenicol plates. Growth on test-plates that contain either ampicillin or no antibiotics served as control for expected loss of the pKD46 plasmid and to make glycerol stocks of bacteria respectively. PCR products with the expected size were sent for sequencing (Eurofins). In order to move the recombineered part of the genome into fresh cells, P1 phage transfer was conducted ^48^.

### *Escherichia coli* ViaA cloning, expression and purification

*E. coli* ViaA (Genebank nr.: AAT48203.1, Uniprot ID: P0ADN0) cloned in the p11-Toronto1 vector and containing an N-terminal hexahistidine (6xHis) tag was obtained from Prof. Dr. Walid Houry. Overlap extension PCR was used to add a C-terminal AviTag (GLNDIFEAQKIEWHE) to the His-ViaA construct. Additionally, Gibson assembly was used to clone ViaA in the pET22b vector containing a C-terminal His-tag and an added N-terminal AviTag.

N- and C-terminally AviTagged ViaA constructs, cloned in the pET22b and p11-Toronto1 vectors respectively, were transformed in chemically competent BL21 (DE3) *E. coli* cells. The resulting transformants were grown in a 50 ml preculture supplemented with Ampicillin to an OD of 3.5. Next, 2 l of LB supplemented with Ampicillin was inoculated with 40 ml of the pre-culture and grown at 37°C until an OD of 0.8 was reached. Cells were subsequently induced with 0.5 mM IPTG and incubated at 18 °C overnight (ON). The next day, the cells were harvested by centrifuging the cultures at 4,000 g for 45 min.

For purification of expressed AviTag-ViaA-His and His-ViaA-AviTag constructs, pellets (4 for a 2 l culture) were resuspended in 30 ml 25 mM HEPES pH 7.5, 300 mM NaCl, 10 mM MgCl2, 10 % Glycerol, 1 μl benzonase, 1 x Complete tablet, and disrupted by three passages through a Homogeniser at 18,5000 psi. Cell debris was removed by centrifugation for 1 h at 20.00048,384g, and the supernatant was filtered using a 0.2 μl filter and flown over a 5 ml NiNTA Immobilised Metal Affinity Chromatography (IMAC) column (GE Healthcare) equilibrated with buffer A (25 mM HEPES pH 7.5, 300 mM NaCl, 10 mM MgCl2, 10 % Glycerol, 25 mM Imidazole). AVI-ViaA-HIS was washed with 5 CV of buffer A, and eluted using a gradient of 0 – 100% buffer B (25 mM HEPES pH 7.5, 300 mM NaCl, 10 mM MgCl2, 10 % Glycerol, 300 mM Imidazole). The top fractions after IMAC were pooled and desalted using a 5 ml HiTrap Desalting column (GE Healthcare) equilibrated with 25 mM HEPES pH 7.5, 300 mM NaCl, 10 mM MgCl2, 10 % Glycerol. After desalting, the buffer exchanged protein was concentrated and injected on a SD200 10/300 increase column (GE Healthcare) equilibrated with 25 mM HEPES pH 7.5, 300 mM NaCl, 10 mM MgCl2, 10 % Glycerol, 1 mM DTT. Purified ViaA fractions, eluted at around 14 ml, were flash-frozen and stored at -80 °C for later use.

### *Escherichia coli* NuoEF, NuoEFG and FrdA expression and purification

Expression of soluble NuoEF and NuoEFG was based on Braun *et al*. ^20^ and Bungert *et al*. ^21^. Chemically competent *E. coli* BL21 (DE3) cells were transformed with a pET-11a vector containing NuoB-G subunits with NuoF harboring an N-terminal Strep-Tag, kindly provided by Thorsten Friedrich ^21^. The resulting transformants were grown for approx. 5 hours in a 2 L culture of LB medium supplemented with 100 mg/L ferric ammonium citrate added in aliquots of 20 mg/L per hour,2 mg/L cysteine, and 20 mg/L riboflavin. Cells were grown at 37 °C and 0.1 mM IPTG was added when the OD reached 0.8. After three hours, cells were pelleted by centrifugation at 4,000g for 10 min. The cell pellet was resuspended in 50 mM MES, 50 mM NaCl, pH 6.6, 10 µg/ml DNase and 1X Complete tablet and disrupted by one passage through a Homogeniser at 18,000 psi. Cell debris was removed by centrifugation for 1 h at 250,000g. The supernatant was filtered using a 0.2 μl filter and flown over a 5 ml StrepTactin-Sepharose column (GE Healthcare) equilibrated with buffer A (50 mM MES-NaOH, 50 mM NaCl, pH 6.6). The column was washed with buffer A, and the NuoEF and EFG fragments were eluted with buffer A supplemented with 2.5 mM D-desthiobiotin. The eluted NuoEF and NuoEFG mixture appeared reddish/brown, and was further purified using a SD200 10/300 increase column (GE Healthcare) equilibrated with 50 mM MES, 500 mM NaCl, pH 6.6, resulting in two separated peaks containing NuoEFG and NuoEF respectively. Fractions containing NuoEFG and NuoEF, verified by SDS-PAGE, were concentrated separately, flash frozen and stored at -80 °C.

Expression of soluble FrdA was based on the protocol by Léger *et al*. ^22^. Chemically competent *E. coli* BL21 (DE3) cells with a knock-out in FrdB were transformed with N-terminal His-tagged FrdA in the pnEA_His6_3C vector, gratefully obtained by Dr. Christophe Romier, Strasbourg. The resulting transformants were used to inoculate 200 ml of LB + 200 µl Ampicillin in a 1 l flask and the culture was grown at 37 °C with vigorous aeration (250 rpm) ON. The pre-culture was used to inoculate 0.5 of LB medium with 500 ml Ampicillin. The resulting culture was grown at 37 °C until the OD reached 0.6. Gene expression was induced with 0.5 mM IPTG. After +-18h of expression at 18 °C, cells were pelleted by centrifugation at 4000 g for 45 min. The cell pellet was resuspended in 30 ml Buffer A (50 mM KH_2_PO_4_ pH 7, 500 mM NaCl, 1% glycerol) supplemented with 1x Complete tablet and 0.5 µl benzonase. Cells were opened by sonication, and cell debris was removed by centrifugation for 1 h at 20,000 g. The supernatant was filtered using a 0.2 μl filter and flown over a 5 ml NiNTA IMAC column (GE Healthcare) equilibrated with buffer A (50 mM KH_2_PO_4_ pH 7, 500 mM NaCl, 1% glycerol). Bound FrdA was washed with 5% buffer B (50 mM KH_2_PO_4_ pH 7, 500 mM NaCl, 1% glycerol, 500 mM Imidazole), followed by elution using a gradient from 5 – 100 % buffer B. Eluted FrdA fractions appeared bright yellow due to the present FAD cofactor. The eluted FrdA fraction was subsequently concentrated and injected onto a SD200 10/300 increase column (GE Healthcare) equilibrated with 20 mM HEPES pH 7.5, 300 mM NaCl, 10% glycerol, 1 mM DTT. Purified FrdA fractions were flash-frozen and stored at -80 °C for later use.

### Multi-Angle Laser light Scattering (MALLS)

Prior to conducting MALLS experiments, flash-frozen purified AviTag-ViaA-His and His-NTV-Avi samples were thawed and centrifuged at 20,000g for 30 min. SEC-MALLS experiments were conducted at 4 °C on a high-performance liquid chromatography (HPLC) system (Schimadzu, Kyoto, Japan) consisting of a DGU**-**20 AD degasser, an LC-20 AD pump, a SIL20-AC_HT_ autosampler, a CBM-20A communication interface, an SPD-M20A UV-Vis detector, a FRC-10A fraction collector, an XL-Therm column oven (WynSep, Sainte Foy d’Aigrefeuille, France) and static light scattering miniDawn Treos, dynamic light scattering DynaPro NANOSTAR and refractive index Optilab rEX detectors (Wyatt, Santa-Barbara, USA).

Purified AviTag-ViaA-His (80 µl at 3 mg/ml) or His-NTV-AviTag (80 µl at 2 mg/ml) were injected on a Superdex 200 increase 10/300 GL column (GE Healthcare), equilibrated at 4°C with a buffer containing 25 mM HEPES pH 7.5, 300 mM NaCl, 10 mM MgCl_2_ and 10 % Glycerol, at a flow rate of 0.5 ml/min. Bovine serum albumin (BSA) at 2 mg/ml in phosphate-buffered saline (PBS) buffer was injected as a control. The extinction coefficient and refractive index increments for the proteins were calculated from the amino acid sequences using the SEDFIT software.

### Small Angle X-Ray Scattering (SAXS)

SAXS data were collected on the BM-29 BIOSAXS beamline at the ESRF (Grenoble, France) ^49^, equipped with a Pilatus3 2M detector operated at a wavelength of 0.9919 Å and using a sample-detector distance 2.867 m, resulting in a scattering momentum transfer range of 0.003 A^-1^ to 0.494 A^-1^. Purified AviTag-ViaA-His was measured at a concentration of 3 mg/ml in a buffer containing 25mM HEPES pH 7.5, 300 mM NaCl, 10 mM MgCl_2_ and 10 % Glycerol. Measurements were performed at 20 °C, and 10 frames with an individual exposure time of 0.5 s were taken per sample. Initial data integration, averaging and background subtraction were performed using PRIMUS-QT from the ATSAS software package ^50^.

The forward scattering (*I*_0_) and radius of gyration (*R*_g_) were determined by PRIMUS-QT using Guinier approximation ^51^. The Porod volume estimate (*V*_p_) was evaluated using Autoporod ^52^. The maximum particle dimension *D*_max_ and distance distribution function *P*(r) were evaluated using ScÅtter ^18^. The molecular mass of the His-ViaA-AviTag sample was calculated using the online SAXSMoW 2.0 program ^17^ and the ScÅtter software package.

### Bio-Layer Interferometry

For BLI binding studies, biotinylated AviTag-ViaA-His, His-ViaA-AviTag and His-NTV-AviTag samples were expressed and purified as described earlier, but with the addition of 50 μM D-biotin to the LB medium during expression overnight at 18 °C. BLI experiments were performed in HBS kinetics buffer, containing 25 mM HEPES pH 7.5, 300 mM NaCl, 10 mM MgCl_2_, 10 % Glycerol, 0.1% w/v BSA and 0.02% v/v Tween-20), using an Octet RED96 instrument (FortéBio) operated at 293 K. Streptavidin-coated biosensors (FortéBio) were functionalised with biotinylated AviTag-ViaA-His, His-ViaA-AviTag or His-NTV-Avi to a maximum signal of 1 nm, subsequently quenched with 10 μg ml^−1^ biocytin, and transferred to wells containing 5 different concentrations of ligand. When RavA or RavAΔLARA was used as a ligand, 1mM ADP was added to the HBS kinetics buffer and RavA RavA or RavAΔLARA samples were incubated with 1 mM ADP for 10 min prior to conducting measurements. Buffer subtraction was performed using a functionalized biosensor measuring running buffer. To check for nonspecific binding during the experiments, non-functionalized biosensors were used to measure the signal from the highest ligand concentration as well as running buffer. All data were fitted with the FortéBio Data Analysis 9.0 software using a 1:1 interaction model. All binding experiments were performed in triplicate (technical triplicates), and calculated K_D_, k_d_ and k_a_ values represent the averages of these triplicate experiments.

### Nanobody production and labelling

The anti-RavA-Nb and anti-ViaA-Nb were obtained from the nanobody generation platform of the AFMB laboratory (Marseille, France) as described ^15^. The anti-BC2-Nb ^23^ was kindly provided by Ulrich Rothbauer. Nanobodies were labelled as described ^15^.

### Overexpression of recombinant RavA and ViaA constructs for fluorescence imaging, cellular EM observations, and lipid extraction and analysis

MG1655 cells were transformed with a low-copy auxiliary pT7pol26 (Kan^R^) plasmid that codes for T7 RNA polymerase under the control of a lac promoter ^53^. The resulting MG1655/pT7pol26 (Kan^R^) strain was then transformed with a plasmid (Amp^R^) carrying a gene coding for a desired RavA or ViaA construct under the control of the T7 promotor. The cells were cultured at 37 °C until the OD_600_ reached 0.6, and then RavA-OE or ViaA-OE was induced with 40 µM IPTG for 12 hours, at 18°C, under aerobic conditions.

### Fluorescence imaging

For immunofluorescence staining cells were harvested, fixed and permeabilised as described ^15^. For epifluorescence imaging of fluorescent protein fusion constructs the procedure was the same except that the fixation was performed with 1% formaldehyde solution in PBS, and that no permeabilisation was necessary.

### Wide field imaging

For each sample, 2 μL of cells in suspension were mounted between a glass slide and a 1.5h glass coverslip, and observed using an inverted IX81 microscope, with the UPLFLN 100× oil immersion objective from Olympus (numerical aperture 1.3), using a fibered Xcite™ Metal-Halide excitation lamp in conjunction with the appropriate excitation filters, dichroic mirrors, and emission filters specific for DAPI/Hoechst, AF488, mCherry or AF647 (4×4MB set, Semrock). Acquisitions were performed with Volocity software (Quorum Technologies) using a sCMOS 2048 × 2048 camera (Hamamatsu ORCA Flash 4, 16 bits/pixel) achieving a final magnification of 64 nm per pixel.

### STORM and PALM imaging

Super-resolution single molecule localization microscopy was performed using STORM (after nanobody labeling) and PALM (PAmCherry fusion proteins) approaches. For STORM imaging, cells labeled as mentioned above, were transferred to a glucose buffer containing 50 mM NaCl, 150 mM Tris (pH 8.0), 10% glucose, 100 mM MEA (mercaptoethylamine) and 1x Glox solution from a 10x stock containing 1 μM catalase and 2.3 μM glucoseoxidase. For PALM, cells expressing the ViaA fused constructs with PAmCherry, were resuspended in PBS. In each case, 2 µL of the cells in suspension were mounted as specified in Jessop and al.(2021). Mounted samples were imaged on an IX81 microscope (Olympus) by focusing the excitation lasers to the back focal plane of an oil immersion UAPON100X (N.A. 1.49) objective. Acquisitions were performed using an Evolve 512 camera with a gain set to 200 (electron-multiplying charge-coupled device EMCCD, 16 bits/pixel, Photometrics) using Metamorph (Molecular Devices). Excitation lasers power was controlled by an Acousto-Optical Tunable Filter (OATF, Quanta Tech). SMLM datasets of about 30,000 frames were generated using 3 kW/cm ^2^ of a 643 nm laser in conjunction with up to 1 W/cm2 of a 405 nm laser at a framerate of 20 frames per seconds (STORM) or approximately 1 kW/cm2 of a 561nm laser after photoactivation of PAmCherry using a few watts/cm ^2^ of a 405 nm laser (PALM). Data processing was performed using Thunderstorm plugin in ImageJ ^54,55^.

### Cellular EM observations

WT, RavA-OE and ViaA-OE cells were centrifuged at 4000 g for 5 min. A pellet volume of 1.4 μl was dispensed on the 200 μm side of a type A 3 mm gold platelet (Leica Microsystems), covered with the flat side of a type B 3 mm aluminum platelet (Leica Microsystems), and was vitrified by high-pressure freezing using an HPM100 system (Leica Microsystems). Next, the samples were freeze substituted at −90 °C for 80 h in acetone supplemented with 1% OsO_4_ and warmed up slowly (1°C h^−1^) to −60 °C in an automated freeze substitution device (AFS2; Leica Microsystems). After 8 to 12 h, the temperature was raised (1°C h^−1^) to -30°C, and the samples were kept at this temperature for another 8 to 12 h before a step for 1 h at 0 °C, cooled down to -30 °C and then rinsed four times in pure acetone. The samples were then infiltrated with gradually increasing concentrations of Epoxy Resin (Epoxy Embedding Medium kit, Merck) in acetone (1:2, 1:1, 2:1 [vol/vol] and pure) for 2 to 8 h while raising the temperature to 20°C. Pure epoxy resin was added at room temperature. After polymerization 24 h at 60°C, 60 to 80 nm sections were obtained using an ultra-microtome UC7 (Leica Microsystems) and an Ultra 35° diamond knife (DiATOME) and were collected on formvar-carbon-coated 100 mesh copper grids (EMS). The thin sections were post-stained for 10 min with 2% uranyl acetate, rinsed and incubated for 5 min with lead citrate. The samples were observed using a FEI Tecnai12 120kV LaB6 microscope with an Orius SC1000 CCD camera (Gatan).

### Lipid extraction

Lipids were extracted from freeze-dried cells. First, cells were harvested by centrifugation and then immediately frozen in liquid nitrogen. Once freeze-dried, the pellet was suspended in 4 mL of boiling ethanol for 5 minutes to prevent lipid degradation and lipids were extracted according to Folch *et al*. ^56^ by addition of 2 mL methanol and 8 mL chloroform. The mixture was saturated with argon and stirred for 1 hour at room temperature. After filtration through glass wool, cell remains were rinsed with 3 mL chloroform/methanol 2:1, v/v and 5 mL of NaCl 1% were added to initiate biphase formation. The chloroform phase was dried under argon and the lipid extract was stored at -20°C.

### TLC and GC-FID/MS phospholipid quantification

Total phospholipids were quantified from their fatty acids. The lipid extract was solubilised in pure chloroform and 5 µg of C21:0 (internal standard) were added in an aliquot fraction. Fatty acids were converted to methyl esters (FAME) by a 1-hour incubation in 3 mL 2.5% H2SO4 in pure methanol at 100°C ^57^. The reaction was stopped by addition of 3 mL water and 3 mL hexane. The hexane phase was analyzed by gas chromatography (Clarus 80, Perkin Elmer) on a BPX70 (SGE) column. FAMEs were identified by comparison of their retention times with those of standards (Sigma) and by their mass fragmentation spectra. They were quantified using C21:0 for calibration. To quantify each class of phospholipid, 300 µg of lipids were separated by two-dimensional thin layer chromatography (TLC) onto glass-backed silica gel plates (Merck) ^58^. The first solvent was chloroform:methanol:water (65:25:4, v/v) and the second one chloroform:acetone:methanol:acetic acid:water (50:20:10:10:5, v/v). Lipids were sprayed with 2% 8-anilino-1-naphthalenesulfonic acid in methanol, then visualised under UV light and scraped off the plate. Lipids in the scraped silica were quantified by methanolysis and GC-FID/MS as described above.

### Dot blot assay (Protein-lipid overlay assay)

All phospholipids were purchased from Avanti Lipids (Sigma). 2 µl of chloroform solutions (5, 0.5 and 0.05 mg/ml) of 1-palmitoyl-2-oleoyl-glycero-3-phosphocholine (16:0-18:1 PC), 1-palmitoyl-2-oleoyl-sn-glycero-3-phospho-L-serine (16:0-18:1 PS), 1-palmitoyl-2-oleoyl-sn-glycero-3-phosphoethanolamine (PE 16:0-18:1 PE), 1-palmitoyl-2-oleoyl-sn-glycero-3-phospho-(1’-rac-glycerol) (16:0-18:1 PG), 1-palmitoyl-2-oleoyl-sn-glycero-3-phosphate (16:0-18:1 PA) and 1’,3’-bis[1-palmitoyl-2-oleoyl-sn-glycero-3-phospho]-glycerol (16:0-18:1 Cardiolipin) were spotted using a Hamilton syringe onto a nitrocellulose membrane (Biorad trans-blot turbo RTA Midi 0.2 µm) to yield 10, 1 and 0.1 µg of the lipid per spot. The membranes were blocked using a Tris-buffered saline (TBS) solution (pH 7.4) supplemented with 10 % (w/v) non-fat milk powder at room temperature for 1 h before incubated with 10 µg/ml of the purified proteins (RavA, ViaA, LdcI) or protein complex (LdcI-RavA) in TBS containing 0.05 % (v/v) Tween-20 (TBST). After overnight incubation at 4 °C, the membranes were washed with TBST (3×15 min) and probed either with anti-ViaA mouse monoclonal antibody or a rabbit polyclonal antibody raised against the purified RavA or LdcI proteins diluted in TBST at room temperature for 1 h. The membranes were washed with TBST (3×15 min) and further incubated with either anti-mouse or anti-rabbit IgG secondary antibody conjugated with horseradish peroxidase (Merk) at room temperature for 1 h. After washing with TBST (3×15 min), the chemiluminescence signal was developed using a Pierce ECL Western blotting substrate (Thermo Fisher Scientific, USA) and recorded using a ChemiDoc XRS+ System (Bio-rad, USA).

For RavA bound via ViaA, the milk-blocked membrane was incubated with TBST alone (RavA only) or with 10µ g/ml ViaA overnight at 4 °C, washed three times with TBST buffer and incubated with 10µ g/ml RavA for 2 h at room temperature. The chemiluminescent signal was developed after the incubation of the membranes with anti-RavA serum followed by incubation with the HRP-conjugated secondary antibody. Likewise, for ViaA bound via RavA, the membrane was incubated with 10µ g/ml RavA overnight before incubation with 10µ g/ml ViaA. The LdcI-RavA complex was prepared by mixing the purified LdcI and RavA proteins in molar ratio of 1:2.25 (LdcI:RavA) before incubating for 30 min at room temperature. The milk-blocked membranes were incubated with RavA (10µ g/ml) or LdcI (6.5 µg/ml) alone or with the LdcI-RavA complex (corresponding to the final concentration of LdcI and RavA proteins of 6.5 µg/ml and 10 µg/ml, respectively) diluted in TBST supplemented with 10 mM MgCl2 and 2 mM ADP. After overnight incubation at 4 °C, the membranes were washed and the chemiluminescent signal was developed after incubation of the membranes with anti-RavA or anti-LdcI sera followed by incubation with the HRP-conjugated secondary antibody (both diluted also in TBST-Mg-ADP buffer).

### Time-dependent killing assay

This experiment was performed as previously described ^5^. Briefly, overnight cultures were diluted (1/100) and grown anaerobically at 37 °C to an OD_600_ of 0.2 in LB medium supplemented with fumarate. At this point (T0), Gm was added to the cells at 16 µg/mL. After 1.5 and 3 hours, 100 μL of cells were diluted in sterile phosphate buffered saline solution (PBS buffer), spotted on LB agar and incubated at 37 °C for 48 hours. Cell survival was determined by counting colony-forming units per mL (CFU/mL). The absolute CFU at time-point zero (used as the 100%) was approximately 5×10^7^ CFU/mL. This assay was performed in an anaerobic chamber (Jacomex, France).

### Planar lipid bilayer experiments

Planar lipid bilayer measurements were performed at 25 °C using a home-made Teflon cells separated by a diaphragm with a circular hole with the diameter of 0.5 mm. The membrane was formed by a painting method using a mixture of lipids composed of PA:PG:PE (10:45:45) dissolved in *n*-decane and 3 % asolectin dissolved in *n*-decane:butanol (9:1) for ViaA and RavA proteins and the *B. pertussis* pore-forming CyaA toxin, respectively. The membrane current was recorded by Theta Ag/AgCl electrodes after addition of 2 nM ViaA and RavA proteins or 250 pM CyaA toxin ^59^ in 10mM Tris-HCl (pH 7.4), 0.15 M KCl and 2 mM CaCl_2_ using applied potential of -50 and +50 mV. The signal was amplified by an LCA-200-100G Ultra-Low-Noise Current Amplifier (Femto, Germany) and digitised by a Keithley KPCI-3108 PCI Bus Data Acquisition Boards card. The signal was analysed using QuB (https://qub.mandelics.com) and processed using a 10-Hz filter.

## Acknowledgements

We thank Alain Roussel and Aline Desmyter for production of anti-RavA and anti-ViaA nanobodies, and Ulrich Rothbauer and Ulrike Endesfelder for sharing their construct of the BC2-tag and their anti-BC2 nanobody with us. We are grateful to Megghane Baulard and Virgile Adam for help with initial construct design and optical imaging, to Beatrice Py for advice and training in generating chromosomal mutants, to Patricia Renesto for initial characterisation of AG sensitivity, to Humaira Khaliq for planar lipid bilayer measurements and to Dominique Bourgeois for discussions. This work was funded by the European Union’s Horizon 2020 research and innovation programme under grant agreement No 647784 to IG. The nanobody generation platform of the AFMB laboratory (Marseille, France) was supported by the French Infrastructure for Integrated Structural Biology (FRISBI) ANR-10-INSB-05-01. We used the platforms of the Grenoble Instruct-ERIC center (ISBG; UAR 3518 CNRS-CEA-UGA-EMBL) within the Grenoble Partnership for Structural Biology (PSB), supported by FRISBI (ANR-10-INBS-0005-02) and GRAL, financed within the University Grenoble Alpes graduate school (Ecoles Universitaires de Recherche) CBH-EUR-GS (ANR-17-EURE-0003). The EM facility is supported by the Rhône-Alpes Region, the Fondation Recherche Medicale (FRM), the fonds FEDER and the GIS-Infrastrutures en Biologie Sante et Agronomie (IBISA). J.F. was supported by a long-term European Molecular Biology Organization (EMBO) fellowship (ALTF441-2017) and a Marie Skłodowska-Curie actions individual fellowship (789385, RespViRALI). L.B. was supported by the project LM2018133 (EATRIS-CZ) of the Ministry of Education, Youth and Sports of the Czech Republic.

## Author contributions

JF, LB, CL, AF, FR, JYEK, KH, BG, CM, JPK, YD, MJ, EK and IG performed experiments, JF, LB, CL, AF, FR, JYEK, JPK, YD, MJ, FB, JJ and IG analysed data. IG designed the overall study, supervised the project and wrote the paper with significant input from JF and contributions from LB, AF, FR, KH, JPK, YD and JJ. JF prepared the figures with contributions from LB, CL, AF, FR, JYEK, JPK, YD and IG. All authors edited the manuscript prior to submission.

## Supplementary Figure Legends

**Supplementary Figure 1:** A) BLI measurements of His-NTV-AviTag coupled on BLI biosensors and RavA, with or without added ADP/ATP. B) BLI measurements of AviTag-ViaA-His coupled on BLI biosensors and RavA_ΔLARA_, with or without added ADP/ATP. For A) and B), the blue curves correspond to the measured signal while the red curves correspond to the calculated fit using a 1:1 (no nucleotide, ADP) or 2:1 heterogeneous ligand binding (ATP) interaction model. C) Molecular mass determination of His-NTV-AviTag by SEC-MALLS. The differential refractive index (RI) signal is plotted (left axis, blue curve) along with the UV signal (UV, left axis, green curve) and the determined molecular weight (MW, right axis, grey curve). The theoretical monomer MW and the MW as determined by MALLS are annotated on the left-hand side of the plot.

**Supplementary Figure 2:** A) Single molecule localisation microscopy imaging of *E. coli* cells overexpressing the N-terminal domain of ViaA (NTV) fused to PAmCherry by PALM at 37° C (left), 30° C (middle) and 18° C (right). B & C) Wide field imaging of *E. coli* cells overexpressing RavA (B) or ViaA (C) using phase contrast, DNA (Hoechst) staining or anti-RavA (VHH2-AF488) or anti-ViaA (VHH1-AF647) nanobodies coupled to Alexa Fluor dyes.

**Supplementary Figure 3:** Dot-blot assays using purified ViaA constructs (A: ViaA, ViaA_1-472_, ViaA_R476E/R479E_), purified RavA constructs (B: RavA, RavA_ΔLARA_), or purified LdcI-RavA complex (C) visualized using anti-ViaA (A) and anti-RavA (B & C) or anti-LdcI (C) antibodies and a secondary HRP-antibody conjugate (PC: phosphatidylcholine, PS: phosphatidylserine, PE: phosphatidyl-ethanolamine, PG: phosphatidylglycerol, PA: phosphatidic acid, CL: cardiolipin).

**Supplementary Figure 4:** TEM imaging of high pressure frozen, freeze-substituted and sectioned *E. coli* cells overexpressing different RavA (blue square: RavA, RavA_ΔLARA_) or ViaA (purple square: ViaA, ViaA_1-472_ or ViaA_R476E/R479E_.) constructs.

**Supplementary Figure 5:** A) Bar chart showing the determination of protein quantity of overexpressed RavA, RavA_ΔLARA_, ViaA, ViaA_1-472_ and ViaA_R476E/R479E_, based on SDS-PAGE gels shown in B (RavA), C (RavA_ΔLARA_), D (ViaA), E (ViaA_1-472_) and F(ViaA_R476E/R479E_).

**Supplementary Figure 6:** A & B) Quantification of fatty acid levels (total, phosphatidylethanolamine, phosphatidylglycerol or cardiolipin) by TLC and GC-FID/MS in wild-type (WT) MG1655 *E. coli* cells and MG1655 *E. coli* cells overexpressing RavA, RavA_ΔLARA_, ViaA, ViaA_R476E/R479E_ and ViaA_1-472_, visualized by bar charts (A) and Tukey representations (B, showing all points from min. to max.). Bars are the average of 3 to 7 measurements (5 for MG1655, 7 for ViaA-OE, 3 for the other mutants), plus or minus SD. The overexpressing (OE) series were compared with the control WT MG1655 series using an unpaired Student t test. In the Tukey representations, each point represents a biological repeat. The overexpressing (OE) series were compared with the control WT MG1655 series using an unpaired nonparametric Mann-Whitney test. For both representations, significant difference with the control is shown by * (p value < 0.05) or ** (p value < 0.01). The # indicates a significant difference (p value < 0.05) with ViaA-OE.

**Supplementary Figure 7:** Overall membrane activities of ViaA, RavA and the *B. pertussi*s pore-forming CyaA toxin as measured by planar lipid bilayer technique. The ViaA and RavA proteins were diluted to a final concentration of 2 nM and exposed to a lipid membrane containing PA, PG and PE lipids in molar ratio of 10:45:45. The CyaA toxin, used as a positive control for membrane pore formation, was diluted to to a final concentration of 250 pM and exposed to an asolectin membrane. The aqueous phase contained 10 mM Tris-HCl (pH 7.4), 150 mM KCl and 2 mM CaCl_2_; the applied voltage was 50 mV; the temperature was 25 °C. The membrane current recordings were processed using a 10 Hz filter.

