## Supplementary information for "The AAA+ ATPase RavA and its binding partner ViaA modulate *E. coli* aminoglycoside sensitivity through interaction with the inner membrane"

**Supplementary Figure and Legends**

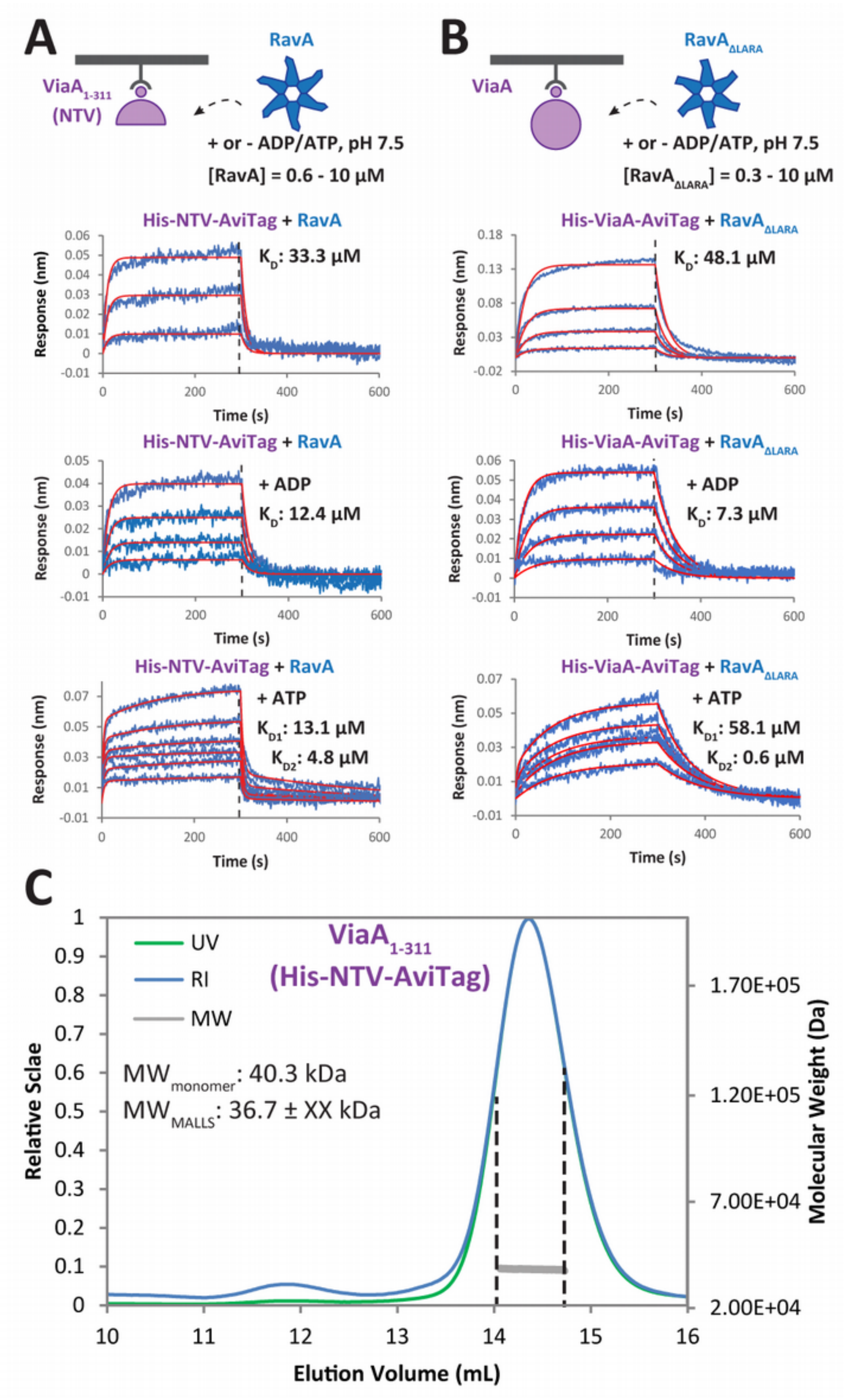


**Supplementary Figure 1:** A) BLI measurements of His-NTV-AviTag coupled on BLI biosensors and RavA, with or without added ADP/ATP. B) BLI measurements of AviTag-ViaA-His coupled on BLI biosensors and $RavA_{\Delta LARA}$, with or without added ADP/ATP. For A) and B), the blue curves correspond to the measured signal while the red curves correspond to the calculated fit using a 1:1 (no nucleotide, ADP) or 2:1 heterogeneous ligand binding (ATP) interaction model. C) Molecular mass determination of His-NTV-AviTag by SEC-MALLS. The differential refractive index (RI) signal is plotted (left axis, blue curve) along with the UV signal (UV, left axis, green curve) and the determined molecular weight (MW, right axis, grey curve). The theoretical monomer MW and the MW as determined by MALLS are annotated on the left-hand side of the plot.

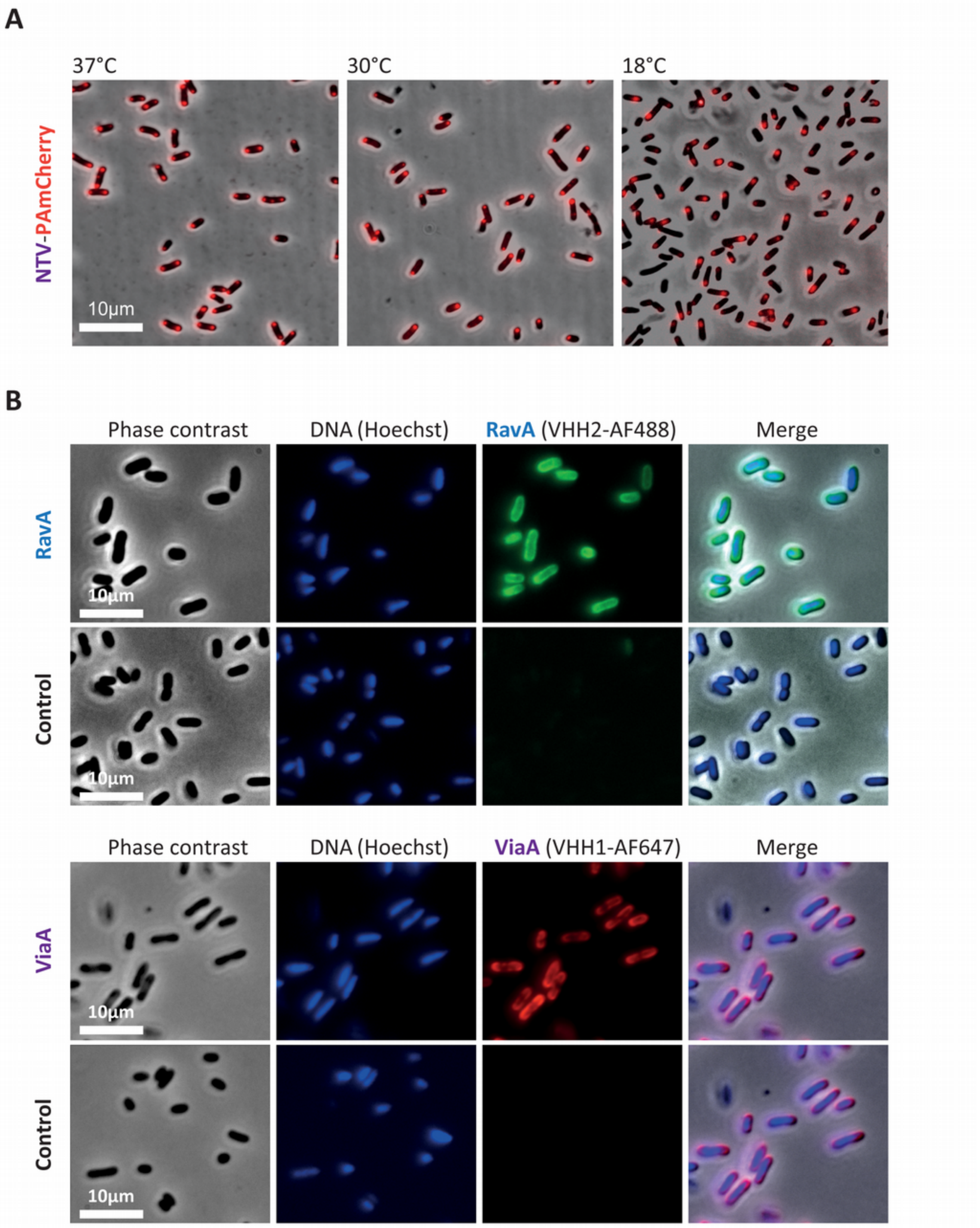


**Supplementary Figure 2:** A) Single molecule localisation microscopy imaging of *E. coli* cells overexpressing the N-terminal domain of ViaA (NTV) fused to PAmCherry by PALM at 37° C (left), 30° C (middle) and 18° C (right). B & C) Wide field imaging of *E. coli* cells overexpressing RavA (B) or ViaA (C) using phase contrast, DNA (Hoechst) staining or anti-RavA (VHH2-AF488) or anti-ViaA (VHH1-AF647) nanobodies coupled to Alexa Fluor dyes.

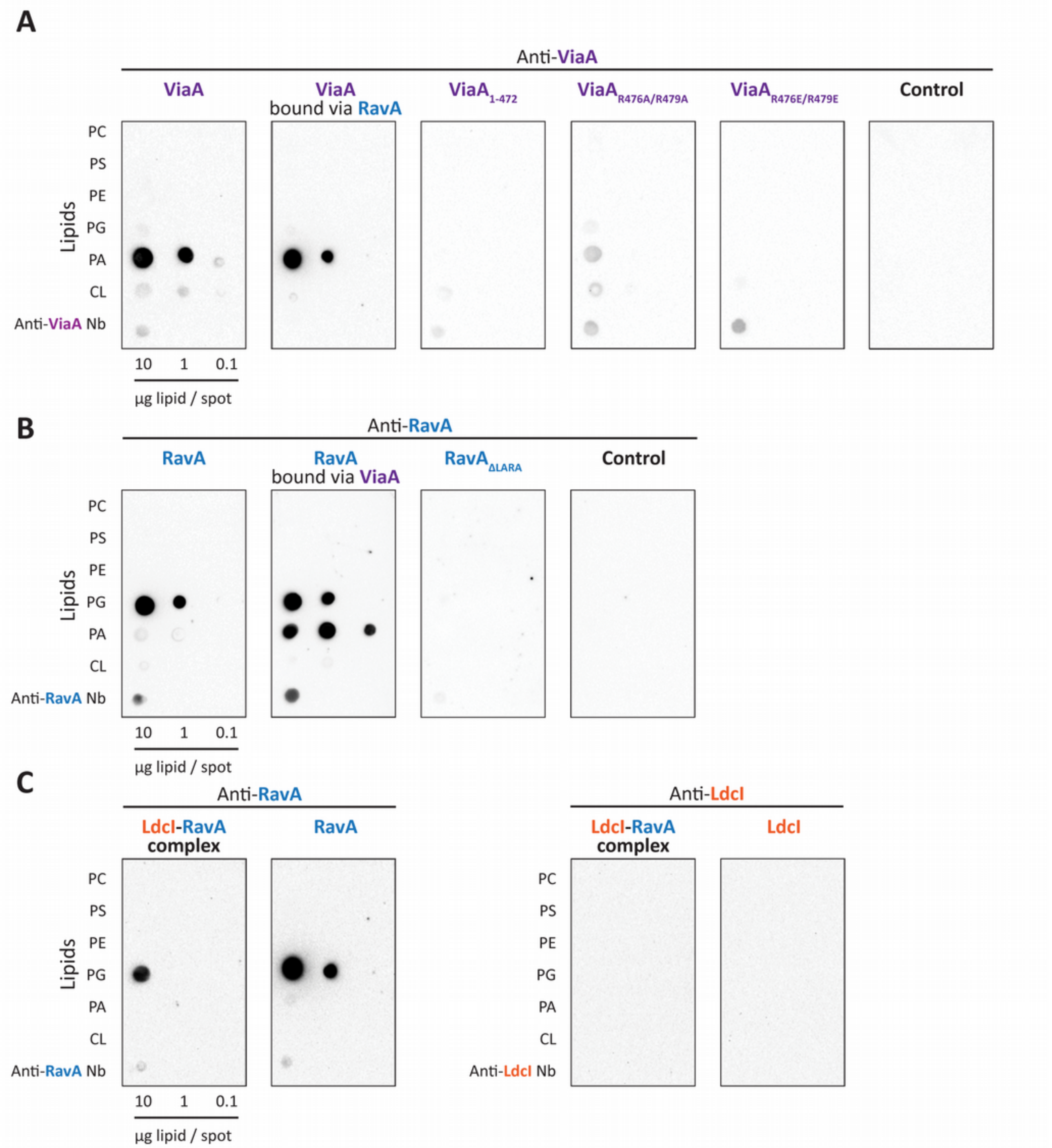


**Supplementary Figure 3:** Dot-blot assays using purified ViaA constructs (A: ViaA, $ViaA_{1\text{-}472}$, $ViaA_{R476E/R479E}$), purified RavA constructs (B: RavA, $RavA_{\Delta LARA}$), or purified LdcI-RavA complex (C) visualized using anti-ViaA (A) and anti-RavA (B & C) or anti-LdcI (C) antibodies and a secondary HRP-antibody conjugate (PC: phosphatidylcholine, PS: phosphatidylserine, PE: phosphatidyl-ethanolamine, PG: phosphatidylglycerol, PA: phosphatidic acid, CL: cardiolipin).

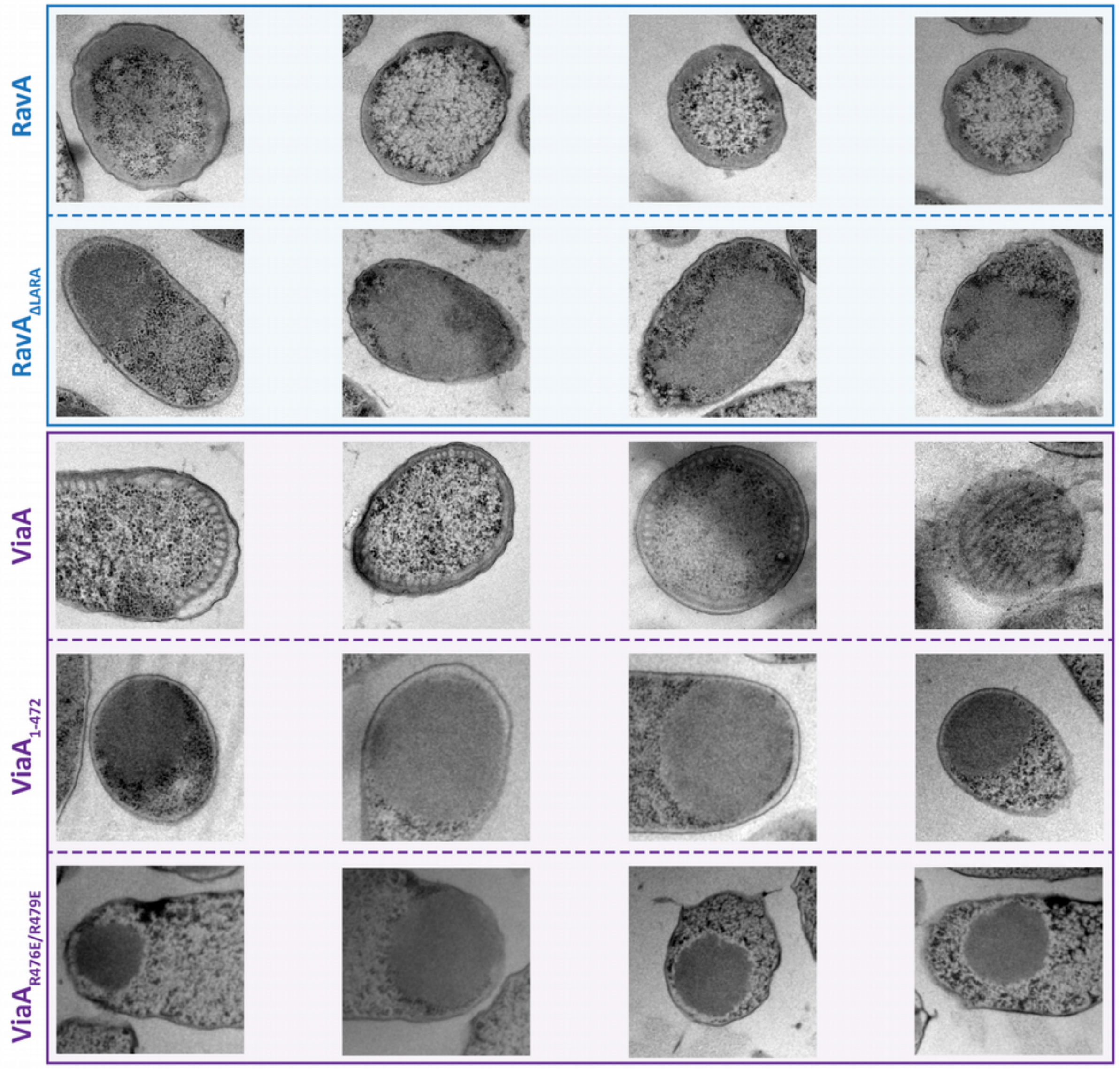


**Supplementary Figure 4:** TEM imaging of high pressure frozen, freeze-substituted and sectioned *E. coli* cells overexpressing different RavA (blue square: RavA, $RavA_{\Delta LARA}$) or ViaA (purple square: ViaA, $ViaA_{1\text{-}472}$ or $ViaA_{R476E/R479E}$.) constructs.

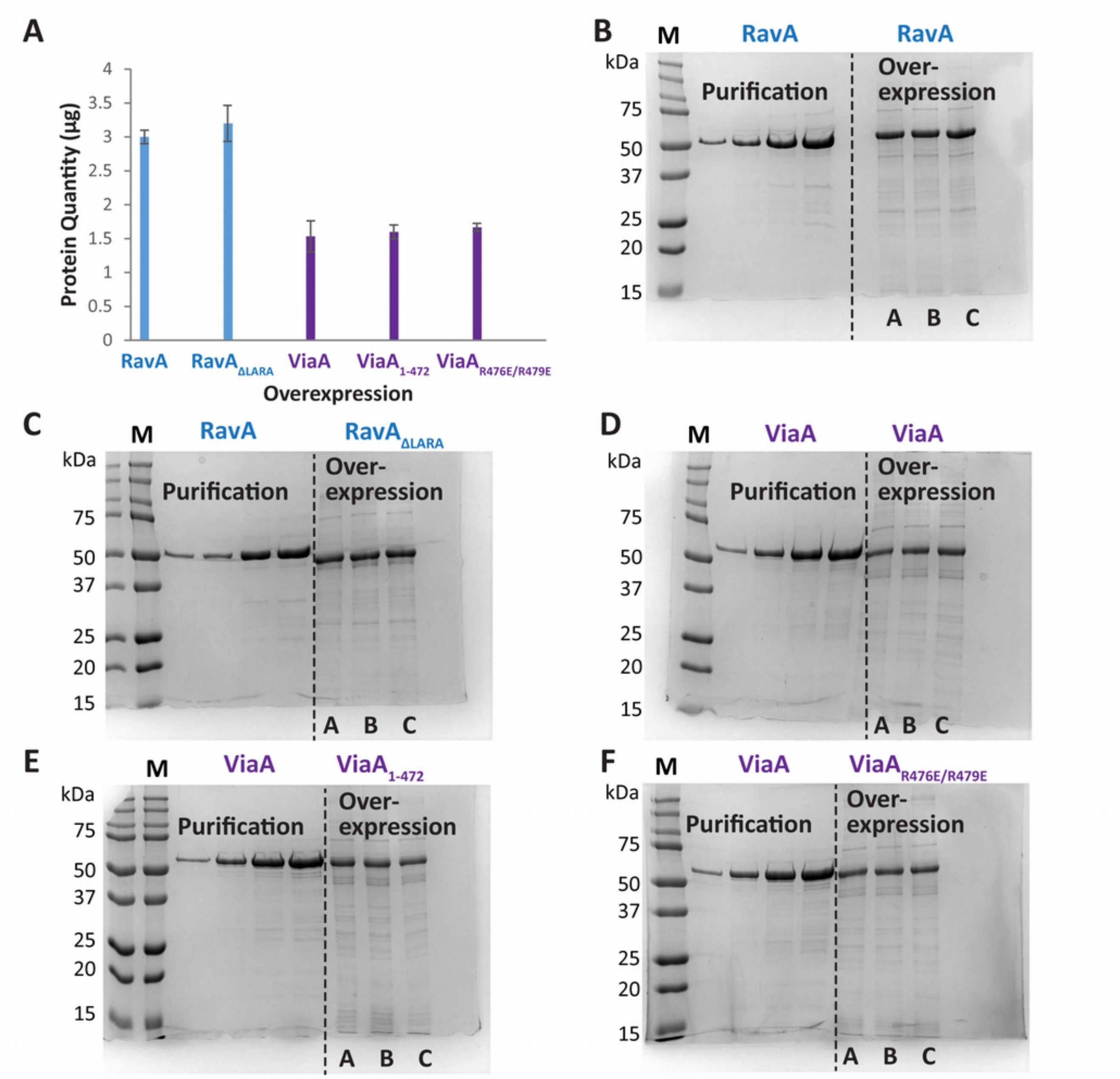


**Supplementary Figure 5:** A) Bar chart showing the determination of protein quantity of overexpressed RavA, $RavA_{\Delta LARA}$, ViaA, $ViaA_{1\text{-}472}$ and $ViaA_{R476E/R479E}$, based on SDS-PAGE gels shown in B (RavA), C ($RavA_{\Delta LARA}$), D (ViaA), E ($ViaA_{1\text{-}472}$) and F($ViaA_{R476E/R479E}$).

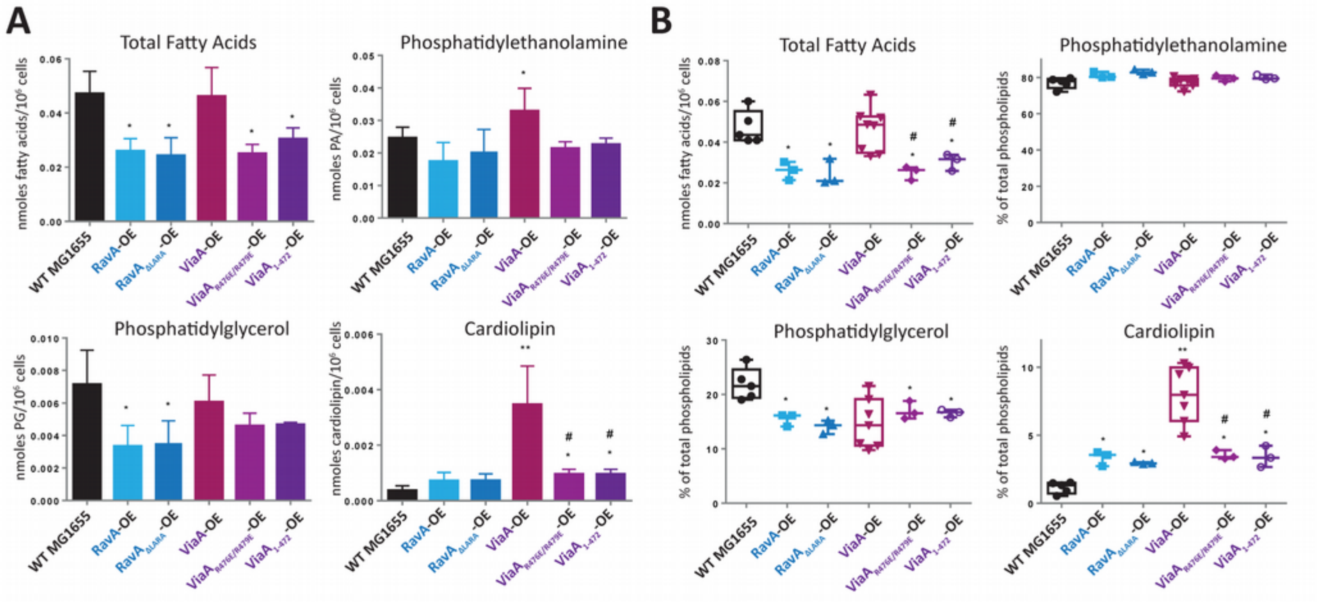


**Supplementary Figure 6:** A & B) Quantification of fatty acid levels (total, phosphatidylethanolamine, phosphatidylglycerol or cardiolipin) by TLC and GC-FID/MS in wild-type (WT) MG1655 *E. coli* cells and MG1655 *E. coli* cells overexpressing RavA, $RavA_{\Delta LARA}$, ViaA, $ViaA_{R476E/R479E}$ and $ViaA_{1-472}$, visualized by bar charts (A) and Tukey representations (B, showing all points from min. to max.). Bars are the average of 3 to 7 measurements (5 for MG1655, 7 for ViaA-OE, 3 for the other mutants), plus or minus SD. The overexpressing (OE) series were compared with the control WT MG1655 series using an unpaired Student t test. In the Tukey representations, each point represents a biological repeat. The overexpressing (OE) series were compared with the control WT MG1655 series using an unpaired nonparametric Mann-Whitney test. For both representations, significant difference with the control is shown by * (p value < 0.05) or ** (p value < 0.01). The # indicates a significant difference (p value < 0.05) with ViaA-OE.

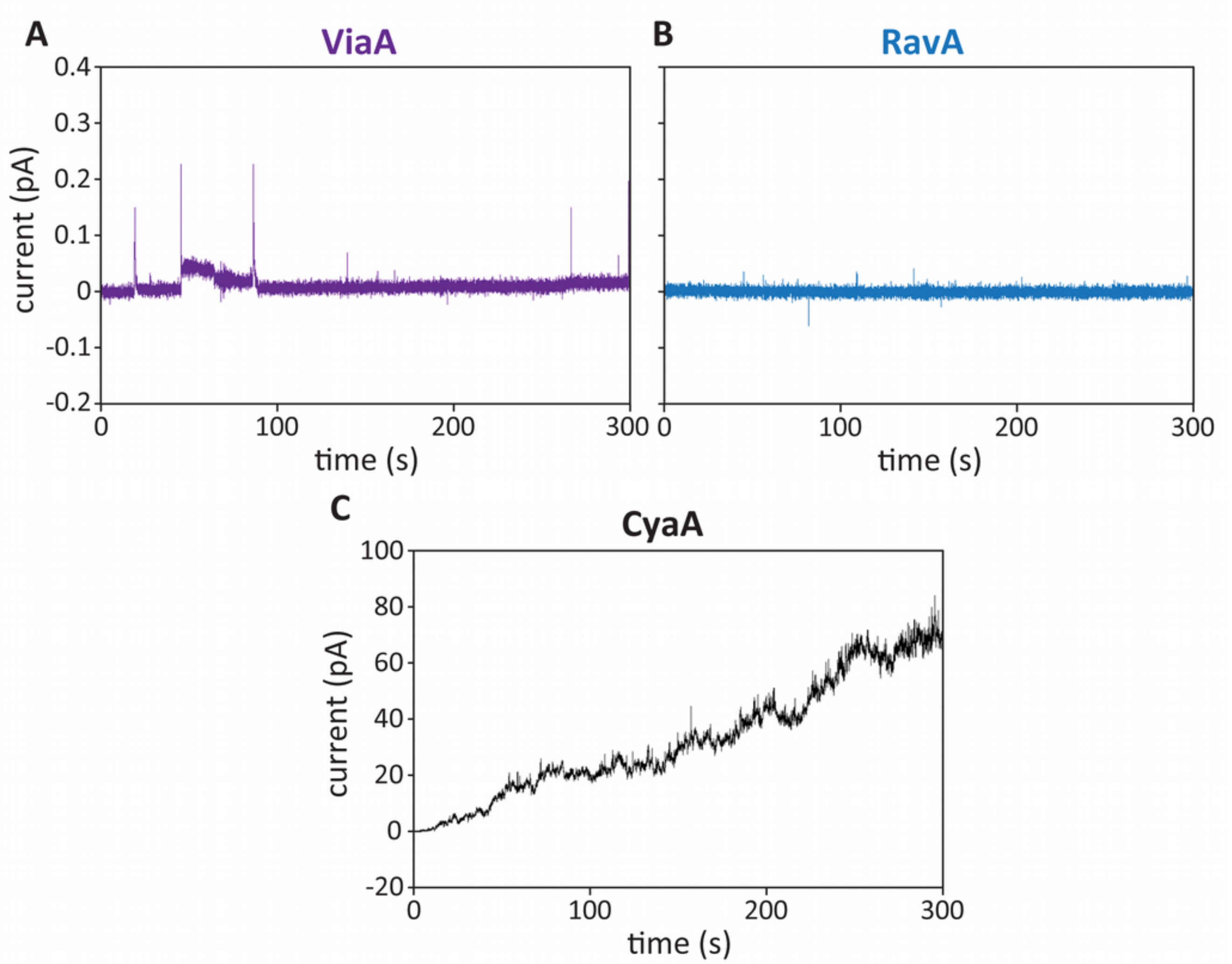


**Supplementary Figure 7:** Overall membrane activities of ViaA, RavA and the *B. pertussis* pore-forming CyaA toxin as measured by planar lipid bilayer technique. The ViaA and RavA proteins were diluted to a final concentration of 2 nM and exposed to a lipid membrane containing PA, PG and PE lipids in molar ratio of 10:45:45. The CyaA toxin, used as a positive control for membrane pore formation, was diluted to to a final concentration of 250 pM and exposed to an asolectin membrane. The aqueous phase contained 10 mM Tris-HCl (pH 7.4), 150 mM KCl and 2 mM $CaCl_2$; the applied voltage was 50 mV; the temperature was 25 °C. The membrane current recordings were processed using a 10 Hz filter.
